# Gene knock-ins in *Drosophila* using homology-independent insertion of universal donor plasmids

**DOI:** 10.1101/639484

**Authors:** Justin A. Bosch, Ryan Colbeth, Jonathan Zirin, Norbert Perrimon

**Affiliations:** Department of Genetics, Harvard Medical School, Boston, MA, 02115, USA; Howard Hughes Medical Institute, Boston, MA, 02115, USA

**Keywords:** CRISPR-Cas9, knock-in, homology-independent, gene function, CRISPaint

## Abstract

Targeted genomic knock-ins are a valuable tool to probe gene function. However, knock-in methods involving homology-directed repair (HDR) can be laborious. Here, we adapt the mammalian CRISPaint homology-independent knock-in method for *Drosophila melanogaster*, which uses CRISPR/Cas9 and non-homologous end joining (NHEJ) to insert universal donor plasmids into the genome. This method is a simple and fast alternative to HDR for certain strategies such as C-terminal tagging and gene disruption. Using this method in cultured S2R+ cells, we efficiently tagged four endogenous proteins with the bright fluorescent protein mNeonGreen, thereby demonstrating that an existing collection of CRISPaint universal donor plasmids is compatible with insect cells. In addition, we inserted the transgenesis marker *3xP3-RFP* into seven genes in the fly germ line, producing heritable loss of function alleles that were isolated by simple fluorescence screening. Unlike in cultured cells, indels always occurred at the genomic insertion site, which prevents predictably matching the insert coding frame to the target gene. Despite this effect, we were able to isolate *T2A-Gal4* insertions in four genes that serve as in vivo expression reporters. Finally, we apply this fast knock-in method to uncharacterized small open reading frame (smORF) genes. Therefore, homology-independent insertion is a useful genome editing technique in *Drosophila* that will better enable researchers to dissect gene function.

**Article summary:** We report a fast and simple genomic knock-in method in *Drosophila* to insert large DNA elements into any target gene. Using CRISPR-Cas9 and non-homologous end joining (NHEJ), an entire donor plasmid is inserted into the genome without the need for homology arms. We demonstrate its usefulness in cultured cells to fluorescently tag endogenous proteins and in the fly germ line to generate heritable insertions that disrupt gene function and can act as expression reporters.

## Introduction

Insertion of DNA into the animal genome is a powerful method to study gene function. This approach is multipurpose, and can be used, for example, to alter gene function, assay gene expression, visualize protein localization, and purify endogenous proteins (Auer and Del bene 2014; Singh *et al.* 2015). Furthermore, the ability to insert large DNA elements such as promoters, protein coding sequences, or entire genes into the genome offers researchers endless options for genome modification. *Drosophila melanogaster* is an excellent animal model to analyze gene function because of its many genetic tools, fast generation time, and in vivo analysis (Venken *et al.* 2016; Korona *et al.* 2017; Bier *et al.* 2018).

Two commonly used methods in *Drosophila* to insert DNA into endogenous genes are transposable DNA elements and homology directed-repair (HDR). Transposable elements insert randomly in the genome (Bellen *et al.* 2011) and thus cannot be used to target a user-specified gene. In contrast, HDR is used to insert DNA into a precise genomic location by homologous recombination (Bier *et al.* 2018). Circular plasmids are commonly used as donor DNA for HDR because they can carry a large DNA insert (≤10kb) and homology arms corresponding to the target locus are added by molecular cloning. However, the design and construction of plasmid donors for each gene can be laborious and prone to troubleshooting. As a cloning-free alternative, synthesized single-stranded DNA (ssDNA) with short homology arms (∼50-100 bp each) (Bier *et al.* 2018) or long ssDNAs of ≤2kb (Quadros *et al.* 2017) can be used as donors. However, ssDNA donors are limited to small insertions, and, like plasmid donors, must be designed and generated for each gene that is targeted by HDR. The burden of generating unique donor DNAs for HDR makes this method inefficient for targeting many genes in parallel, such as hits from a genetic or proteomic screen, or the same gene in divergent species. Therefore, there is a need for simpler and more scalable alternatives to knock-in large DNA elements into the *Drosophila* genome.

Large DNA elements can also be inserted in target genes without homology arms, known as homology-independent insertion (Cristea *et al.* 2013; Maresca *et al.* 2013; Auer *et al.* 2014; Katic *et al.* 2015; Lackner *et al.* 2015; Schmid-Burgk *et al.* 2016; Suzuki *et al.* 2016; Katoh *et al.* 2017; Kumagai *et al.* 2017; Zhang *et al.* 2018; Gao *et al.* 2019). In this method, simultaneous cutting of a circular donor plasmid and a genomic target-site by a nuclease results in integration of linearized insert DNA into the genomic cut site by non-homologous end joining (NHEJ). Such donor plasmids are “universal” because they lack gene-specific homology arms, allowing insert sequences (e.g. GFP) to be targeted to any genomic location. Despite this advantage, homology-independent insertion is less precise than HDR. For example, donor DNA can insert in two directions, insertion and deletion mutations (indels) at the integration site can affect the insert translation frame, the entire plasmid can integrate, and inserts can form concatemers. Nevertheless, for some targeting strategies such as C-terminal protein tagging or gene disruption, homology-independent insertion is a fast, simple, and effective alternative to HDR. While homology-independent insertion has been successfully applied in human cell lines (Cristea *et al.* 2013; Maresca *et al.* 2013; Lackner *et al.* 2015; Schmid-Burgk *et al.* 2016; Katoh *et al.* 2017; Zhang *et al.* 2018), mouse somatic cells (Suzuki *et al.* 2016; Gao *et al.* 2019), zebrafish (Auer *et al.* 2014), *C. elegans* (Katic *et al.* 2015), and *Daphnia* (Kumagai *et al.* 2017), it has not yet been utilized in *Drosophila*.

Here, we use the CRISPaint strategy to show that homology-independent insertion functions effectively in *Drosophila*. First, we inserted an mNeonGreen universal donor plasmid into the C-terminus of four target proteins in S2R+ cells and used Puromycin-resistance to select for in-frame insertions, demonstrating that this is an efficient method to produce cell lines with tagged endogenous proteins. Second, we inserted a *T2A-Gal4 3xP3-RFP* universal donor plasmid into the fly germ line and isolated knock-in lines for seven genes by simple fluorescent screening. By targeting insertions to 5’ coding sequence, we demonstrate that this is an effective method to disrupt gene function and generate loss of function fly strains. Furthermore, by screening insertions for Gal4 expression, we identified four that are in vivo reporters of their target gene. Finally, we use this knock-in method to quickly generate insertions in uncharacterized small open reading frame (smORF) genes.

## Materials and Methods

### Plasmid cloning

*pCFD3-frame_selector_*(*0, 1, or 2*) plasmids (Addgene #127553-127555, DGRC #1482-1484) were cloned by ligating annealed oligos encoding sgRNAs that target the CRISPaint target site (Schmid-Burgk *et al.* 2016) into *pCFD3* (Port *et al.* 2014), which contains the *Drosophila U6:3* promoter.

Additional sgRNA-encoding plasmids were generated by the TRiP (https://fgr.hms.harvard.edu/) or obtained from Filip Port (Port *et al.* 2015). sgRNA plasmids targeting CDS close to the stop codon were *GP07595* (*Act5c*) (Addgene #130278, DGRC #1492), *GP07596* (*His2Av*), *GP07609* (*alphaTub84B*), and *GP07612* (*Lam*). sgRNA plasmids targeting CDS close to the start codon were *GP06461* (*wg*), *GP02894* (*FK506-bp2*), *GP05054* (*alphaTub84B*), *GP00225* (*esg*), *GP00364* (*Myo1a*), *GP00400* (*btl*), *GP00583* (*Mhc*), *GP01881* (*hh*), *GP03252* (*Desat1*), *GP05302* (*ap*), *pFP545* (*ebony*), and *pFP573* (*ebony*). These sgRNAs were cloned into *pCFD3*, with the exception of those targeting *esg, Myo1a, btl*, and *Mhc*, which were cloned into *pl100* (Kondo and Ueda 2013).

*pCRISPaint-T2A-Gal4-3xP3-RFP* (Addgene #127556, DGRC #1481) was constructed using Gibson assembly (E2611, NEB) of three DNA fragments: 1) *Gal4-SV40-3xP3-RFP* was PCR amplified from *pHD-Gal4-DsRed* (Xu et al. 2019 in preparation, (Gratz *et al.* 2014); 2) linear plasmid backbone generated by digesting *pWalium10-roe* (Perkins *et al.* 2015) with AscI/SacI; and 3) a synthesized double-stranded DNA fragment (gBlock, IDT) encoding the CRISPaint target site, linker sequence, T2A, and ends that overlap the other two fragments.

*pCRISPaint-3xP3-RFP* (Addgene # 130276, DGRC #1490) was constructed by digesting *pCRISPaint-T2A-Gal4-3xP3-RFP* with Not1/BamHI to remove *T2A-Gal4* sequences and using Gibson to add a gBlock with a multiple cloning site for future manipulation.

*pCRISPaint-3xP3-GFP* (Addgene # 130277, DGRC #1491) was constructed by Gibson assembly of PCR-amplified backbone from *pCRISPaint-3xP3-RFP* and PCR-amplified EGFP sequence from pM37 (Lee *et al.* 2018).

*pCRISPaint-T2A-ORF-3xP3-RFP* donor plasmids (Addgene #127557-127565, DGRC #1485-1489,1493-1497) were cloned by PCR amplifying the ORFs and Gibson cloning into *CRISPaint-T2A-Gal4-3xP3-RFP* cut with NheI/KpnI. ORF sequences were amplified from templates as follows: *sfGFP* [amplified from *pUAS-TransTimer* (He *et al.* 2019)], *LexGAD* [amplified from p*CoinFLP-LexGAD/Gal4* (Bosch *et al.* 2015)], *QF2* amplified from Addgene #80274, *Cas9-T2A-GFP* (amplified from template kindly provided by Raghuvir Viswanatha, Cas9 from *Streptococcus pyogenes*), *FLPo* (amplified from Addgene #24357), *Gal80* (amplified from Addgene #17748), *Nluc* (amplified from Addgene #62057), *Gal4DBD*, (amplified from Addgene #26233), and *p65* (amplified from Addgene #26234).

*pCRISPaint-sfGFP-3xP3-RFP* (Addgene #127566, DGRC #1486) was cloned by PCR amplifying *sfGFP* coding sequence and Gibson cloning into *CRISPaint-T2A-Gal4-3xP3-RFP* cut with NotI/KpnI.

*pCFD5-frame_selectors_0,1,2* (Addgene #131152) was cloned by PCR amplifying two fragments encoding the three sgRNAs that cut the CRISPaint target site and Gibson cloning into *pCFD5* (Port and Bullock 2016). This plasmid allows expression of sgRNAs that are separated by tRNA sequences, and thus *pCFD5-frame_selectors_0,1,2* expresses all three frame-selector sgRNAs simultaneously.

*pLHA-T2A-Gal4-3xP3-RFP-RHA_ebony* was cloned by digesting *pCRISPaint-T2A-Gal4-3xP3-RFP* with AscI/SacI and purifying insert and backbone DNA together (QIAQuick, Qiagen). Homology arms were amplified from *Drosophila* genomic DNA (*yw*, single fly) using Phusion polymerase (LHA: 1592bp, RHA: 1544bp). Insert, backbone, and both homology arms were assembled by Gibson cloning.

See Supplemental Table 7 for oligo and dsDNA sequences and Addgene and DGRC for plasmid sequences.

### Cell culture

*Drosophila* S2R+ cells stably expressing Cas9 and a mCherry protein trap in *Clic* (known as PT5/Cas9) (Viswanatha *et al.* 2018) were cultured at 25°C using Schneider’s media (21720-024, ThermoFisher) with 10% FBS (A3912, Sigma) and 50 U/ml penicillin strep (15070-063, ThermoFisher). S2R+ cells were transfected using Effectene (301427, Qiagen) following the manufacturer’s instructions. Plasmid mixes were composed of sgRNA-expressing plasmids (see above) and *pCRISPaint-mNeon-PuroR* (Schmid-Burgk *et al.* 2016). Cells were transfected with plasmid mixes in 6-well dishes at 1.8×10^6^ cells/ml, split at a dilution of 1:6 after 3-4 days, and incubated with 2 µg/ml Puromycin (540411 Calbiochem). Every 3-5 days, the media was replaced with fresh Puromycin until the cultures became confluent (∼12-16 days). For single-cell cloning experiments, cultures were split 1:3 two days before sorting. Cells were resuspended in fresh media, triturated to break up cell clumps, and pipetted into a cell straining FACS tube (352235 Corning). Single cells expressing mNeonGreen were sorted into single wells of a 96 well plate containing 50% conditioned media 50% fresh media using an Aria-594 instrument at the Harvard Medical School Division of Immunology’s Flow Cytometry Facility. Once colonies were visible by eye (3-4 weeks), they were expanded and screened for mNeonGreen fluorescence.

### Fly genetics and embryo injections

Flies were maintained on standard fly food at 25°C. Fly stocks were obtained from the Perrimon lab collection or Bloomington Stock center (indicated with BL#). Stocks used in this study are as follows: *yw* (Perrimon Lab), *yw/Y hs-hid* (BL8846), *yw; nos-Cas9attP40/CyO* (derived from BL78781), *yw;; nos-Cas9attP2* (derived from BL78782), *yw; Sp hs-hid/CyO* (derived from BL7757), *yw;; Dr hs-hid/TM3,Sb* (derived from BL7758), *UAS-2xGFP* (BL6874), *wg1-17/CyO* (BL2980), *wg1-8/CyO* (BL5351), *Df(2L)BSC291/CyO* (BL23676), *Mhc[k10423]/CyO* (BL10995), Df(2L)H20/CyO (BL3180), *Df(2L)ED8142/SM6a* (BL24135), *hh[AC]/TM3 Sb* (BL1749), *Df(3R)ED5296/TM3, Sb* (BL9338), *esgG66/CyO UAS-GFP* (BL67748), *Df(2R)Exel6069/CyO* (BL7551), ywCre; D/TM3, Sb (BL851), Dp(2;1)Sco[rv23]; Df(2L)Sco[rv23], b[1] pr[1]/CyO (BL6230).

For embryo injections, each plasmid was column purified (Qiagen) twice, eluted in injection buffer (100 µM NaPO4, 5 mM KCl), and adjusted to 200ng/µl. Plasmids were mixed equally by volume, and mixes were injected into *Drosophila* embryos using standard procedures. For targeting genes on Chr. 2, plasmid mixes were injected into *yw;; nos-Cas9attP2* embryos. For targeting genes on Chr. 3, plasmid mixes were injected into *yw; nos-Cas9attP40/CyO* embryos. Approximately 500 embryos were injected for each targeted gene.

Injected G0 flies were crossed with *yw*. We used *yw/Y hs-hid* to facilitate collecting large numbers of virgin flies by incubating larvae and pupae at 37°C for 1hr. G1 flies were screened for RFP expression in the adult eye on a Zeiss Stemi SVII fluorescence microscope. G1 RFP+ flies were crossed with the appropriate balancer stock (*yw; Sp hs-hid/CyO* or *yw;; Dr hs-hid/TM3,Sb)*. G2 RFP+ males that were *yellow*- (to remove the *nos-Cas9* transgene) and balancer+ were crossed to virgins of the appropriate balancer stock (*yw; Sp hs-hid/CyO* or *yw;; Dr hs-hid/TM3,Sb*). G3 larvae and pupae were heat shocked at 37°C for 1hr to eliminate the *hs-hid* chromosome, which generates a balanced stock (e.g. yw; [RFP+]/CyO).

### Imaging

S2R+ cells expressing mNeonGreen were plated into wells of a glass-bottom 384 well plate (6007558, PerkinElmer). For fixed cell images, cells were incubated with 4% paraformaldehyde for 30min, washed with PBS with .1% TritonX-100 (PBT) 3x 5min each, stained with 1:1000 DAPI (D1306, ThermoFisher) and 1:1000 phalloidin-TRITC (P1951, Sigma), and washed with PBS. Plates were imaged on an IN Cell Analyzer 6000 (GE) using a 20x or 60x objective. Time-lapse videos of live mNeonGreen expressing single cell cloned lines were obtained by taking an image every minute using a 60x objective. Images were processed using Fiji software.

Wing imaginal discs from 3^rd^ instar larvae were dissected in PBS, fixed in 4% paraformaldehyde, and permeabilized in PBT. For Wg staining, carcasses were blocked for 1hr in 5% normal goat serum (S-1000, Vector Labs) at room temp, and incubated with 1:50 mouse anti-wg (4D4, DSHB) primary antibody and 1:500 anti-mouse 488 (A-21202, Molecular Probes) secondary antibody. Primary and secondary antibody incubations were performed at 4°C overnight. All carcasses were stained with DAPI and phalloidin-TRITC, and mounted on glass slides with vectashield (H-1000, Vector Laboratories Inc.) under a coverslip. Images of mounted wing discs were acquired on a Zeiss 780 confocal microscope.

Larvae, pupae, and adult flies were imaged using a Zeiss Axio Zoom V16 fluorescence microscope.

### Quantification of mNeonGreen expressing S2R+ cells

For FACs-based cell counting, we collected cultures from each gene knock-in experiment before and after puromycin selection. Pre-selection cultures were obtained by collecting of 500ul of culture 3-4 days after transfection. Post-selection cultures were obtained after at least 2 weeks of puromycin incubation. Non-transfected cells were used as a negative control. 100,000 cells were counted for each sample and FlowJo software was used to analyze and graph the data. FSC-A vs GFP-A was plotted and we defined mNeonGreen+ cells by setting a signal intensity threshold where <0.02% of negative controls are counted due to autofluorescence.

For microscopy-based cell counting, the number of mNeonGreen cells was quantified by analyzing confocal images in Fiji using the manual Cell Counter Plugin (model). For transfected cells, 6 fields containing at least 200 cells were quantified (i.e. n=6). For puro-selected cells, 3 fields containing at least 200 cells were quantified (i.e. n=3).

### Western blotting

Single cell-cloned cell lines were grown until confluent and 1ml of resuspended cells was centrifuged at 250g for 10min. The cell pellet was resuspended in 1ml ice cold PBS, re-centrifuged, and the pellet was lysed in 250ul 2x SDS-Sample buffer and boiled for 5min. 10ul was loaded on a 4-20% Mini-Protean TGX SDS-Page gel (4561096, BioRad), transferred to PVDF membrane (IPFL00010, Millipore), blocked in 5% non-fat dry milk, primary blotting using mouse anti-mNeonGreen (1:1000, Chromtek 32F6) or hFAB™ Rhodamine Anti-Actin (12004164 BioRad), and secondary blotting using 1:3000 anti-mouse HRP (NXA931, Amersham), imaging using ECL (34580, ThermoFisher) on a ChemiDoc MP Imaging System (BioRad).

### PCR, sequencing, and sgRNA cutting assays

S2R+ cell genomic DNA was isolated using QuickExtract (QE09050, Lucigen). Fly genomic DNA was isolated by grinding a single fly in 50µl squishing buffer (10 mM Tris-Cl pH 8.2, 1 mM EDTA, 25 mM NaCl) with 200µg/ml Proteinase K (3115879001, Roche), incubating at 37°C for 30 min, and 95°C for 2 minutes. PCR was performed using Taq polymerase (TAKR001C, ClonTech) when running DNA fragments on a gel, and Phusion polymerase (M-0530, NEB) was used when DNA fragments were sequenced. DNA fragments corresponding to *mNeonGreen* or *T2A-Gal4* insertion sites were amplified using primer pairs where one primer binds to genomic sequence and the other primer binds to the insert. For amplifying non-knock-in sites, we used primers that flank the sgRNA target site. Primer pairs used for gel analysis and/or Sanger sequencing were designed to produce DNA fragments <1kb. Primer pairs used for next-generation sequencing of the insertion site were designed to produce DNA fragments 200-280bp. DNA fragments were run on a 1% agarose gel for imaging or purified on QIAquick columns (28115, Qiagen) for sequencing analysis. See Supplemental Table 7 for oligo sequences.

Sanger sequencing was performed at the DF/HCC DNA Resource Core facility and chromatograms were analyzed using Lasergene 13 software (DNASTAR). Next-generation sequencing was performed at the MGH CCIB DNA Core. Fastq files were analyzed using CRISPresso2 (Clement *et al.* 2019) by entering the PCR fragment sequence into the exon specification window and setting the window size to 10 bases. Quantification of insertion types (seamless, in-frame indel, and frameshift indel) was taken from the allele plot and frameshift analysis outputs of CRISPresso2. The small proportion of “unmodified” reads that were not called by frameshift analysis were not included in the quantification.

T7 endonuclease assays (M0302L, NEB) were performed following the manufacturer instructions.

Splinkerette sequencing was performed as previously described (Potter and Luo 2010). Briefly, genomic DNA was isolated by single fly squishing (described above) and digested using enzyme BstYI (R0523S, NEB) at 60°C overnight and heat inactivated for 20min at 80°C. Digested DNA was ligated with annealed splinkerette oligonucleotides overnight at 16°C. PCR annealing temperatures were 60°C (round 1) and 64°C (round 2). Round 2 PCR products were either run on an agarose gel and gel fragments purified using QIAquick Gel Extraction Kit (28704, Qiagen), or purified using ExoSAP-IT PCR clean up reagent (78200.200.UL, Applied Biosystems). Oligo sequences used for splinkerette sequencing are listed in Supplemental Table 7. 3’ splinkerette sequence traces, using the *M13R* sequencing primer, frequently mapped to *vermillion*, even when genomic DNA was isolated from yw flies. We believe this resulted from contamination, since commonly used plasmids and fly lines (e.g. *pCFD3* and *pValium20*) have *M13R* sequence adjacent to the *vermillion+* marker transgene.

### Data availability

Plasmids generated in this paper will be available from Addgene and the Drosophila Genomics Resource Center (DGRC). Gal4-expressing fly strains will be available at the Bloomington Drosophila Stock Center (BDSC). Two mNeonGreen-expressing S2R+ cell lines are available at the DGRC (His2Av-mNeonGreen clone B11, DGRC #295; Lamin-mNeonGreen clone D6, DGRC #296). The remaining flies, cells, and sequence data are available from the Perrimon lab on request. Oligo and dsDNA sequences are listed in Supplemental Table 7.

## Results

To test if homology-independent insertion works in *Drosophila*, we implemented a strategy known as CRISPaint (Schmid-Burgk *et al.* 2016). This system is used to insert sequence encoding a protein tag or reporter gene into the coding sequence of an endogenous gene. Although it was originally developed for mammalian cell culture, CRISPaint has several advantages for use in *Drosophila*. First, this system uses CRISPR-Cas9 to induce double-strand breaks (DSBs), which is known to function efficiently in *Drosophila* cultured cells (Bottcher *et al.* 2014; Viswanatha *et al.* 2018) and the germ line (Kondo and Ueda 2013; Ren *et al.* 2013; Yu *et al.* 2013; Bassett *et al.* 2014). Second, its use of a modular frame-selector system makes it simple to obtain insertions that are translated with the target gene. Third, a collection of existing CRISPaint donor plasmids (Schmid-Burgk *et al.* 2016) containing common tags (e.g. GFP, RFP, Luciferase) are seemingly compatible for expression in *Drosophila*.

The CRISPaint system works by introducing three components into Cas9-expressing cells: 1) a single guide RNA (sgRNA) targeting a genomic locus; 2) a donor plasmid containing an insert sequence; and 3) a frame selector sgRNA targeting the donor plasmid (Figure 1A). This causes cleavage of both the genomic locus ansd the donor plasmid, leading to the integration of the entire linearized donor plasmid into the genomic cut site by non-homologous end joining (NHEJ). This insertion site destroys both sgRNA target sites and will no longer be cut. While indels can occur at the insertion site due to error-prone NHEJ, a remarkably high percentage of insertions in human cells are seamless (48-86%) (Schmid-Burgk *et al.* 2016). Therefore, users have some control over their insertion frame by using one of three frame-selector sgRNAs. Importantly, these frame-selector sgRNAs do not target the *Drosophila* genome.

**Figure 1.**
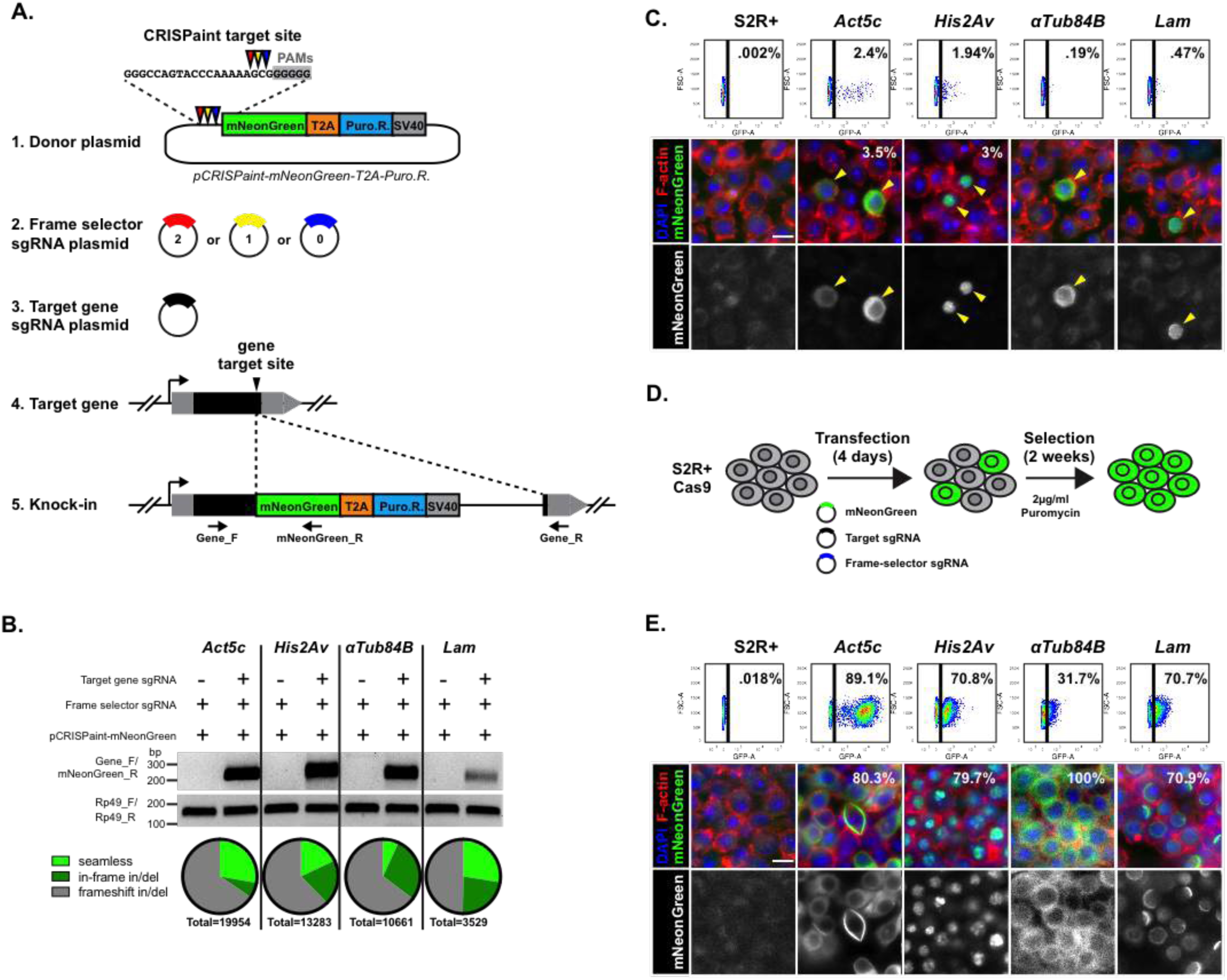
Knock-in of *mNeonGreen-T2A-PuroR* into Drosophila S2R+ cells using homology-independent insertion. (**A**) Schematic of CRISPaint knock-in approach. *mNeonGreen-T2A-PuroR* is inserted into 3’ coding sequence. (**B**) Analysis of knock-in efficiency of transfected cells by diagnostic PCR (DNA gel image) and next-generation sequencing (pie charts). (**C**) Analysis of knock-in efficiency of transfected cells by FACs and confocal microscopy. Numbers indicate percentage of cells with fluorescence. F-actin stained using Phalloidin-TRITC (red), nuclei labeled with DAPI (blue), mNeonGreen signal is in green. Scale bar 10µm. (**D**) Schematic of Puromycin selection of mNeonGreen-expressing cells. (**E**) Analysis of knock-in frequency of puromycin-selected cells using FACs and confocal microscopy. Numbers indicate percentage of cells with green fluorescence. Scale bar 10µm.

### Homology-independent insertion functions efficiently in *Drosophila* S2R+ cells to tag endogenous proteins

To test the CRISPaint method in *Drosophila*, we set out to replicate the findings of (Schmid-Burgk *et al.* 2016) in cultured S2R+ cells by tagging endogenous proteins at their C-terminus. To accomplish this, we generated plasmids expressing frame-selector sgRNAs (frame 0,1, or 2) under the control of *Drosophila U6* sequences (Port *et al.* 2015) (Figure 1A). In addition, we generated plasmids expressing sgRNAs that target the 3’ coding sequence of *Drosophila* genes. We chose to target *Actin5c, His2Av, alphaTub84B*, and *Lamin* because these genes are expressed in S2R+ cells (Hu *et al.* 2017) and encode proteins with known subcellular localization (actin filaments, chromatin, microtubules, nuclear envelope, respectively). For donor plasmid, we used *pCRISPaint-mNeonGreen-T2A-PuroR* (Schmid-Burgk *et al.* 2016), which contains a frame-selector sgRNA target site upstream of coding sequence for the fluorescent mNeonGreen protein and Puromycin resistance protein (PuroR) linked by a cleavable T2A peptide sequence. Only integration of the donor plasmid in-frame with the target coding sequence should result in translation of mNeonGreen-T2A-PuroR (Figure 1A).

We transfected Cas9-expressing S2R+ cells (Viswanatha *et al.* 2018) with a mix of three plasmids: *pCRISPaint-mNeonGreen-T2A-PuroR* donor, target-gene sgRNA, and the appropriate frame-selector sgRNA (Figure 1A, Supplemental Table 1). As an initial method to detect knock-in events, we used PCR to amplify the predicted insertion sites from transfected cells. Using primers that are specific to the target gene and *mNeonGreen* sequence, we successfully amplified sense-orientation *gene-mNeonGreen* DNA fragments for all four genes (Figure 1B). Furthermore, next-generation sequencing of these amplified fragments revealed that 34-50% of sense-orientation insertions are in frame with the target gene (Figure 1B, Supplemental Figure 1, Supplemental Table 1). 7-27% of sense-orientation insertions are seamless, which is slightly lower than previously observed (Schmid-Burgk *et al.* 2016).

Next, we quantified in-frame knock-in frequency by measuring mNeonGreen fluorescence in transfected S2R+ cells. Flow cytometry-based cell counting of transfected cells revealed that the number of mNeonGreen+ cells range from 0.19-2.4% (Figure 1C, Supplemental Table 1), in agreement with human cultured cells (Schmid-Burgk *et al.* 2016). These results were confirmed by confocal analysis of transfected cells, which showed mNeonGreen fluorescence in a small subset of cells (Figure 1C). 3.2% (*Act5c*) and 2.4% (*His2Av*) of transfected cells expressed mNeonGreen (Figure 1C, Supplemental Table 1), which roughly agreed with flow cytometry cell counting. Finally, mNeonGreen localized to the expected subcellular compartments, most obviously observed by His2Av-mNeonGreen and Lam-mNeonGreen co-localization with the nucleus, and Act5c-mNeonGreen and alphatub-mNeonGreen exclusion from the nucleus (Figure 1C). These results suggest that a significant number of transfected S2R+ cells received in-frame insertion of mNeonGreen at their C-terminus using the CRISPaint homology-independent insertion method.

For knock-in cells to be useful in experiments, it is important to derive cultures where most cells, if not all, carry the insertion. Therefore, we enriched for in-frame insertion events using Puromycin selection (Figure 1D). After a two-week incubation of transfected S2R+ cells with Puromycin, flow-cytometry and confocal analysis revealed that 31.7-89.1% of cells expressed mNeonGreen and exhibited correct subcellular localization (Figure 1E, Supplemental Table 1). For *alphaTub84B*, cell counting by flow-cytometry greatly underestimated the number of mNeonGreen+ cells counted by confocal analysis, likely because mNeonGreen expression level was so low. These results demonstrate that Puromycin selection is a fast and efficient method of selecting for mNeonGreen expressing knock-in cells.

A subset of cells in Puro-selected cultures had no mNeonGreen expression or unexpected localization (Figure 1E). Since each culture is composed of different cells with independent insertion events, we used FACs to derive single-cell cloned lines expressing mNeonGreen for further characterization (Figure 2A). At least 14 single-cell cloned lines were isolated for each target gene and imaged by confocal microscopy. Within a given clonal culture, each cell exhibited the same mNeonGreen localization (Figure 2B), confirming our single-cell cloning approach and demonstrating that the insertion is genetically stable over many cell divisions. Importantly, while many clones exhibited the predicted mNeonGreen localization, a subset of the clonal cell lines displayed an unusual localization pattern (Figure 2B). For example, 3/21 Act5c-mNeonGreen clones had localization in prominent rod structures, and 12/14 Lamin-mNeonGreen clones had asymmetric localization in the nuclear envelope (Figure 2B). In addition, some clones had diffuse mNeonGreen localization in the cytoplasm and nucleus (Figure 2B).

**Figure 2.**
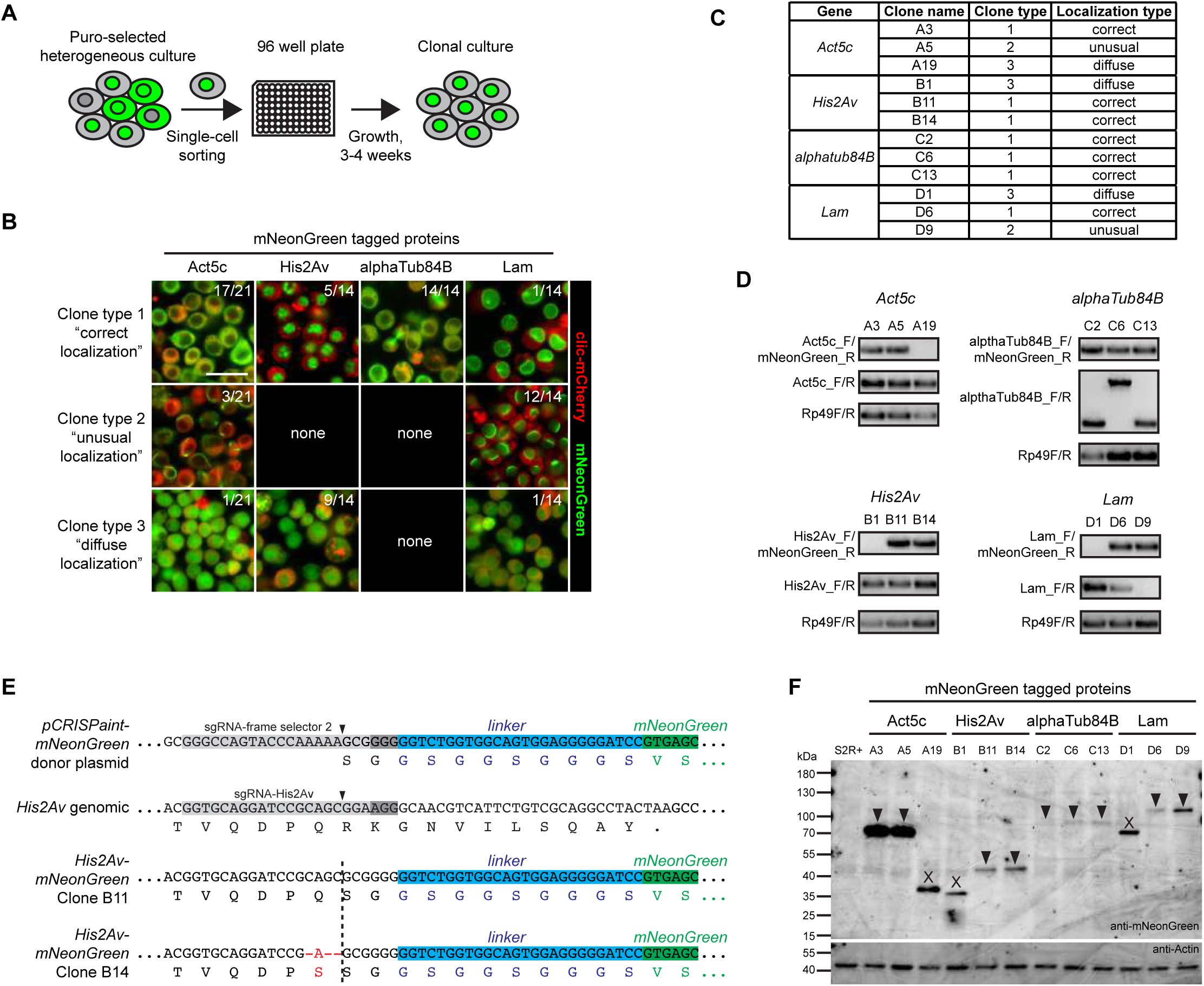
Analysis of S2R+ mNeonGreen-expressing single-cell cloned lines. (**A**) Schematic of FACS isolation of single-cell clones expressing mNeonGreen. (**B**) Confocal images of live mNeonGreen-expressing cell lines, categorized into three clone types. Numbers indicate the frequency of each clone type for each gene targeted. Images show fluorescence from Clic-mCherry (red) and mNeonGreen (green). Scale bar 25µm. (**C**) Single cell cloned lines retained for further analysis. (**D**) Agarose gel with PCR fragments amplified from knock-in (Gene_F/mNeonGreen_R) and non-knock-in loci (Gene_F/R). Positive control bands were amplified from *Rp49* genomic sequence. (**E**) Example sequence results of *His2Av-mNeonGreen* clones. sgRNA target site and PAM sequence shown as grey bars, Cas9 cut sites shown with arrowheads. (**F**) Western blot detecting mNeonGreen protein fusions. Arrowheads indicate expected molecular weight. X’s indicate incorrect molecular weight.

To better characterize the insertions in single cell-cloned lines, we further analyzed three clones per gene (12 total), selecting different classes when possible (correct localization, unusual localization, and diffuse localization) (Figure 2C, Supplemental Table 2). Using PCR amplification of the predicted insertion site (Figure 1A, Figure 2D) and sequencing of amplified fragments (Figure 2E, Supplemental Table 2), we determined that all clones with correct or unusual mNeonGreen localization contained an in-frame insertion of mNeonGreen with the target gene. In contrast, we were unable to amplify DNA fragments from the expected insertion site in clones with diffuse mNeonGreen localization (Figure 2D). Western blotting of cell lysates confirmed that only clones with in-frame mNeonGreen insertion express fusion proteins that match the predicted molecular weights (Figure 2F). All together, these results suggest that clones with correct mNeonGreen localization are likely to contain an in-frame insert in the correct target gene.

S2R+ cells are polyploid (Lee *et al.* 2014), and clones expressing mNeonGreen could bear one or more insertions. Furthermore, indels induced at the non-insertion locus could disrupt protein function. To explore these possibilities, we amplified the non-insertion locus in our single-cell cloned lines and used Sanger and next-generation sequencing to analyze the DNA fragments (Figure 2D, Supplemental Table 2). For each gene, we could find indels occurring at the non-insertion sgRNA cut site. For example, we could distinguish four distinct alleles in clone B11: a 3bp deletion, a 2bp deletion, a 1bp deletion, and a 27bp deletion. In addition, we identified an unusual mutation in clone C6, where a 1482bp DNA fragment inserted at the sgRNA cut site, which corresponds to a region from *alphatub84D*. We assume that this large insertion was caused by homologous recombination, since *alphatub84D* and *alphatub84B* share 92% genomic sequence identity (FlyBase, http://flybase.org/). For *Act5c-mNeonGreen* clones A5 and A19, numerous indel sequences were found, suggesting this region has an abnormal number of gene copies. We were unable to amplify a DNA fragment from *Lam-mNeonGreen* D9, despite follow-up PCRs using primers that bind genomic sequence further away from the insertion site (not shown).

### Loss of function knock-in fly lines by homology-independent insertion in the germ line

We next explored whether homology-independent insertion could function in the *Drosophila* germ line for the purpose of generating knock-in fly strains. As opposed to antibiotic selection in cultured cells, we wanted to identify transgenic flies using a visible body marker that was not dependent on target gene expression. In addition, we wanted to target insertions to 5’ coding sequence to create loss of function alleles by premature protein truncation. Finally, we wanted to test if insertion of a reporter could capture the expression of the target gene. To accomplish these goals, we constructed a new universal donor plasmid called *pCRISPaint-T2A-Gal4-3xP3-RFP* (Figure 3A). This donor contains a frame-selector sgRNA target site upstream of *T2A-Gal4*, which encodes a self-cleaving form of the transcription factor (Diao and White 2012). This donor plasmid also contains the transgenesis marker *3xP3-RFP*, which expresses red fluorescence in larval tissues and the adult eye (Berghammer *et al.* 1999) (Figure 3A).

**Figure 3.**
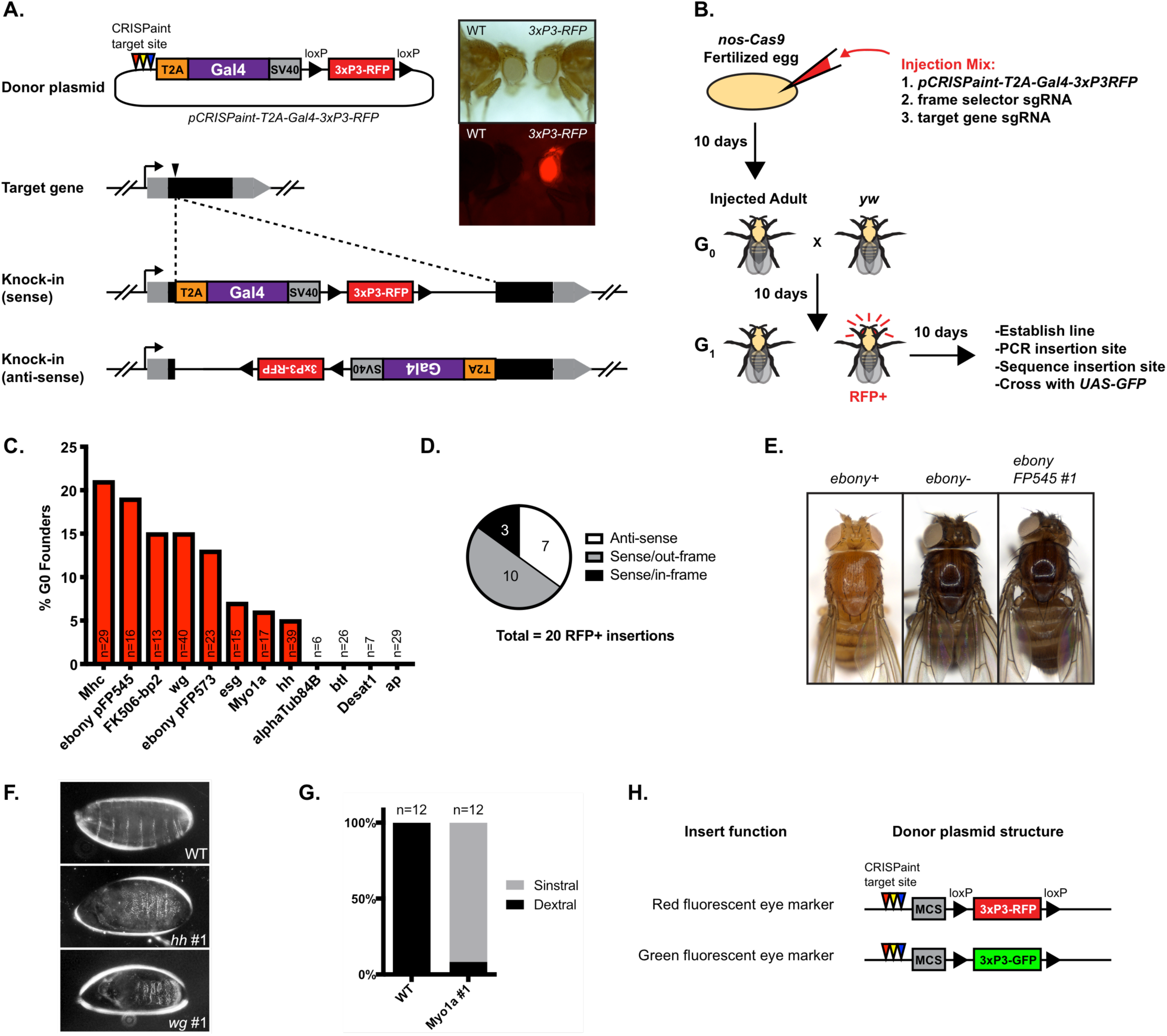
Germ line knock-ins using homology-independent insertion. (**A**) Schematic of knock-in approach. *pCRISPaint-T2A-Gal4-3xP3-RFP* is inserted into 5’ coding sequence. Inset shows images of adult flies with 3xP3-RFP fluorescence in the eye. Top panel is brightfield, bottom panel is fluorescence. (**B**) Schematic of plasmid injections, fly crosses, and analysis of insertions. (**C**) Graph with results of knock-in efficiency for 12 sgRNA target sites and 11 genes. (**D**) Insert orientation and frame of 20 RFP+ fly lines. (**E**) Image of adult flies. Homozygous *ebony-T2A-Gal4 FP545* #1 flies have dark cuticle pigment. (**F**) Darkfield images of *hh* #1 and *wg* #1 homozygous embryo “lawn of denticles” phenotypes. (**G**) Quantification of genital rotation in wild type (*yw*) and a *myo1a* insertion line. Rotation is determined from the direction of wrapping of the adult male spermiduct around the gut from dissected live abdomens. (**H**). Diagram of two simplified universal donor plasmids for germ line knock-ins.

To test if homology-independent insertion functions in the germ line and determine its approximate efficiency, we targeted 11 genes using *pCRISPaint-T2A-Gal4-3xP3-RFP* (Supplemental Table 3). These genes were selected based on their known loss of function phenotypes, expression patterns, and availability of sgRNA plasmids from the TRiP (https://fgr.hms.harvard.edu/). Plasmid mixes of donor, frame-selector sgRNA, and target gene sgRNA were injected into *nos-Cas9* embryos, and the resulting G0 progeny were crossed to *yw*. G1 progeny were screened for RFP fluorescence in adult eyes, and each RFP+ founder fly was crossed to balancer flies to establish a stable stock. Figure 3C and Supplemental Table 3 shows the integration efficiency results for each gene and Supplemental Table 4 has information on 20 RFP+ lines, each derived from a different G0 founder. For injections that produced RFP+ animals, the frequency of G0 crosses yielding RFP+ G1 progeny ranges from 5-21% (Figure 3C, Supplemental Table 3). For example, when targeting *ebony* with sgRNA *pFP545*, 3 out of 16 G0 crosses produced ≥1 RFP+ G1 flies. In one month following embryo injection, we obtained balanced RFP+ lines for seven out of 11 genes targeted (64%) (Figure 3C, Supplemental Table 3).

Next, we used PCR and sequencing to confirm the insertion sites in our RFP+ lines. For each target site, we used two primer pairs to detect the presence of an on-target insertion as well as its orientation (Supplemental Figure 2). Gel images of PCR amplified DNA showed that all 20 of our RFP+ lines contained an insertion in the correct target site (Supplemental Table 4, Supplemental Figure 2), where 13 insertions were in the sense orientation and seven were antisense (Figure 3D). Sequence analysis of PCR fragments confirmed that the insertions were present at the target site (Supplemental Table 4, Supplemental Figure 3). Unexpectedly, all 20 lines contained genomic sequence indels at the insertion site (Supplemental Figure 3), unlike the frequent seamless insertions observed in cultured cells. Germ line indels were also predominantly longer than indels in cultured cells. In particular, a remarkably long 1896bp genomic deletion was found at the *hh* insertion site. Regardless, these results demonstrate that all 20 independently derived RFP+ lines contained *pCRISPaint-T2A-Gal4-3xP3-RFP* at the target site.

Next, we analyzed insertion lines for loss of function phenotypes. Flies with insertions in *wg, Mhc, hh*, and *esg* were homozygous lethal (Supplemental Table 4), which is consistent with characterized mutations in these genes (FlyBase). Similarly, complementation tests revealed that insertions in *wg, Mhc, hh*, and *esg* were lethal in trans with loss of function alleles or genomic deletions spanning the target gene (Supplemental Table 4). In addition, homozygous insertions in *hh* and *wg* cause a “lawn of denticles” phenotype in embryos (Bejsovec and Martinez Arias 1991) (Figure 3F) and flies with heterozygous insertions in *Mhc* were flightless (Mogami *et al.* 1986) (data not shown), all reminiscent of classical allele phenotypes. Four insertions in *ebony* produced flies with dark cuticle pigment when homozygous (Figure 3E, Supplemental Table 4) or in trans with *ebony*^*1*^ (not shown). In one case (*ebony pFP545 #2)*, the insertion was homozygous lethal, but was viable over *ebony*^*1*^ and trans-het flies exhibited dark cuticle pigment. An insertion in *myo1a* (also known as *Myo31DF*) was viable and flies exhibited genital rotation phenotypes observed in previously reported loss-of-function alleles (Speder *et al.* 2006) (Figure 3G). An insertion in *FK506-bp2* was homozygous viable, though this gene has not been well characterized in *Drosophila*. In all, these results indicate that insertion of *pCRISPaint-T2A-Gal4-3xP3-RFP* into 5’ coding sequence can disrupt gene function.

An important consideration when using genome editing is off-target effects. For example, in addition to on-target insertions, our RFP+ fly lines could contain off-target insertions or other mutations that disrupt non-target genes. We noticed that RFP fluorescence always co-segregated with the target chromosome when balancing insertions (not shown), suggesting that multiple insertions may be uncommon. Furthermore, we did not detect evidence of off-target insertions by splinkerette PCR (Potter and Luo 2010), which largely reconfirmed the on-target sites (Supplemental Table 4). However, this analysis was problematic because some sequence traces aligned to the donor plasmid or to genomic sequence on the opposite expected side of the target site, suggesting that insertions could be concatemers. Others have observed concatemers when performing NHEJ knock-ins (Auer *et al.* 2014; Kumagai *et al.* 2017), and we found that seven of our RFP+ insertions were head-to-tail concatemers as determined by PCR (Supplemental Figure 4). We could rescue the lethality of our *esg* insertion using a duplication chromosome (stock # BL6230), suggesting there were no other second site lethal mutations on this chromosome. Finally, one of five *ebony* insertions (*ebony pFP545 #2*) was homozygous lethal, suggesting it contains a second site lethal mutation.

**Figure 4.**
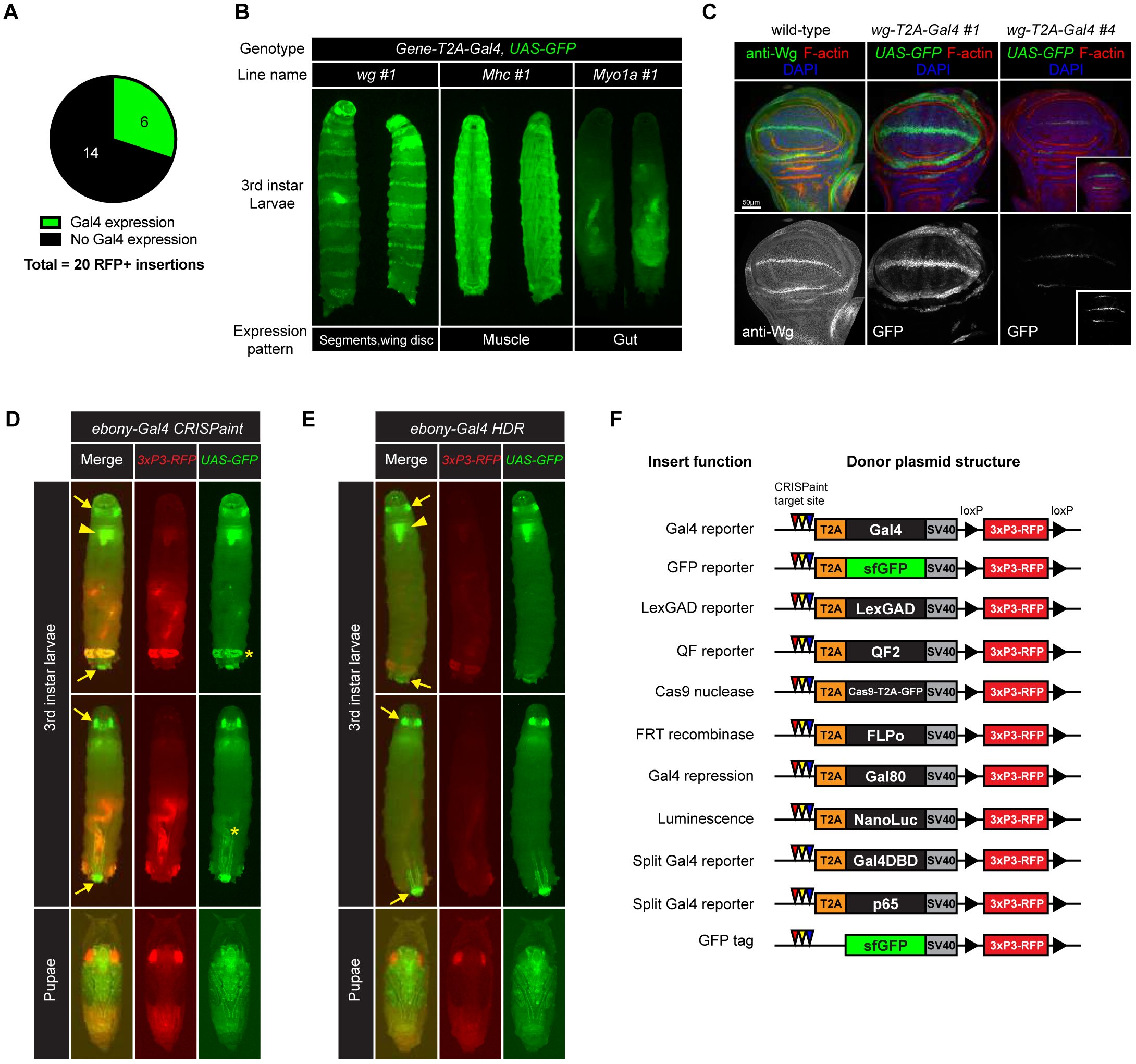
Germ line insertions that express *T2A-Gal4* under the control of the target gene. (**A**) Proportion of 20 RFP+ knock-in lines that express Gal4. (**B**) Fluorescence images of 3^rd^ instar larvae with indicated genotypes. Two larvae of the same genotype are shown as rotated in different angles. Expression of Gal4 under control of the target gene drives expression of the *UAS-GFP* reporter. (**C**) Confocal images of wing imaginal discs showing protein staining of Wg protein (anti-wg, green) or *UAS-GFP* expression (green). GFP fluorescence was recorded at identical exposure settings for lines *wg-T2A-Gal4* #1 and #4. Inset shows digitally increased GFP signal. Scale bar 50µm. (**D**,**E**) Fluorescence images of 3rd instar larvae and pupae. Fluorescence is from *3xP3-RFP* (red) and Gal4 expression with *UAS-GFP* (green). (**D**) shows results with *ebony-T2A-Gal4 pFP545* #2 (*ebony-Gal4* CRISPaint) and (**E**) shows results with *ebony-Gal4* HDR. (**F**) Additional universal donor plasmids for insertion into 5’ coding sequence.

We did not obtain insertions when targeting *ap, alphaTub84B, btl*, or *Desat1*. Therefore, we investigated whether the sgRNAs targeting these genes were functional. 4 sgRNAs used for germ line knock-ins have an acceptable efficiency score of >5, with the exception being the sgRNA targeting *btl* (Supplemental Table 5). A T7 endonuclease assay from transfected cells revealed that sgRNAs targeting *ap, alphaTub84B, btl* can cut at the target site, whereas the results with *desat1* were inconclusive (Supplemental Figure 5A). As an alternative functional test, we used PCR to detect knock-in events in S2R+ cells transfected with the *pCRISPaint-T2A-Gal4-3xP3-RFP* donor plasmid. This showed that sgRNAs targeting *ap, alphaTub84B, btl* and *desat1* can lead to successful knock-in of *pCRISPaint-T2A-Gal4-3xP3-RFP* and suggests that the sgRNAs are functional (Supplemental Figure 5B). Finally, we sequenced the sgRNA target sites in the *nos-Cas9* fly strains and found a SNP in the *btl* sgRNA binding site (not shown), whereas all 10 remaining sgRNAs had no SNPs in the target site. In summary, we conclude that the sgRNAs targeting *ap, alphaTub84B, btl*, and *Desat1* are able to induce cleavage at their target site in S2R+ cells, but that the sgRNA targeting *btl* may not function in the germ line using our *nos-Cas9* strains.

Collectively, our results demonstrate that the *pCRISPaint-T2A-Gal4-3xP3-RFP* universal donor plasmid inserts into target loci in the fly germ line and can be used to generate loss of function fly lines. Therefore, we generated two simplified universal donor plasmids that were more tailored to this purpose, *pCRISPaint-3xP3-RFP* and *pCRISPaint-3xP3-GFP* (Figure 3H). These donor plasmids contain green or red fluorescent markers and a useful multiple cloning site for adding additional insert sequence.

### Knock-ins that express Gal4 under the control of the target gene

We next determined if any of our 20 RFP+ lines expressed Gal4 from the target gene. Since the *T2A-Gal4* reporter gene in *pCRISPaint-T2A-Gal4-3xP3-RFP* is promoterless, we reasoned that only insertions in the sense orientation and in-frame with the target gene should express Gal4. Nonetheless, we took an unbiased approach by crossing all 20 RFP+ lines to *UAS-GFP* and screening progeny for GFP fluorescence throughout development. From this effort, we identified 6 Gal4-expressing lines (Figure 4A, Supplemental Tables 3,4). *wg-T2A-Gal4* (#1 and 4), *Mhc-T2A-Gal4* (#1 and 2), and *Myo1a-T2A-Gal4* #1 insertions are expressed in the imaginal disc, larval muscle, and larval gut (Figure 4B, Supplemental Table 4), respectively, which matches the known expression patterns for these genes. *wg-T2A-Gal4 #1 and #4* insertions are expressed in a distinctive Wg pattern in the wing disc pouch (Figure 4C). *ebony-T2A-Gal4 pFP545 #2* is expressed in the larval brain and throughout the pupal body, consistent with a previous study (Hovemann *et al.* 1998). Interestingly, it is also expressed in the larval trachea, with particularly strong expression in the anterior and posterior spiracles (Figure 4D). Indeed, classical *ebony* mutations are known to cause dark pigment in larval spiracles (Brehme 1941).

Next, we compared the Gal4 expression screening results with our sequence analysis of the insertion sites. As expected, all lines that expressed Gal4 had insertions that were in the sense orientation (Supplemental Table 4, Supplemental Figure 2). For example, *ebony-T2A-Gal4 pFP545 #2* contains a 15bp genomic deletion that is predicted to keep *T2A-Gal4* in-frame with *ebony* coding sequence. Similarly, *wg-T2A-Gal4* #1 contains an in-frame 45bp deletion and 21bp insertion. Remarkably, *wg-T2A-Gal4* #4 contains a frameshift indel (Supplemental Figure 3), yet still expresses Gal4 in the Wg pattern, albeit at significantly lower levels than *wg-T2A-Gal4* #1 (Figure 4C). In addition, *Mhc-T2A-Gal4* lines #1, #2, and *Myo1a-T2A-Gal4* #1, each have indels that put *T2A-Gal4* out of frame with the target gene coding sequence. These findings confirm that our Gal4-expressing lines have *T2A-Gal4* inserted in the sense orientation, but that in-frame insertion with the target gene coding sequence is not necessarily a requirement for Gal4 expression.

Artifacts due to homology-independent insertion, such as indels at the insertion site, could conceivably interfere with Gal4 expression. For example, *ebony* expression has not been well studied, so we did not know if *ebony-T2A-Gal4 pFP545 #*2 (“*ebony-Gal4* CRISPaint”) was an accurate reporter. To address this issue, we generated a precise in-frame HDR insertion in *ebony* (“*ebony-Gal4* HDR”) using the same insert sequence (*T2A-Gal4-3xP3-RFP*) and target gene sgRNA (*pFP545*). *ebony-Gal4* HDR knock-in fly lines were validated by PCR (Supplemental Figure 6) and *ebony-Gal4* HDR homozygous flies were viable and had dark cuticle pigment (not shown), suggesting on-target knock-in. By crossing CRISPaint and HDR knock-in alleles with *UAS-GFP*, we found that their expression patterns were similar at larval, pupal, and adult stages (Figure 4D,E). However, *ebony-Gal4* CRISPaint had higher levels of RFP fluorescence and expressed Gal4 in the larval anal pad and gut, which is coincident with expression of *3xP3-RFP*. Since this insertion is a concatemer (Supplemental Figure 4, Supplemental Table 4), we speculated that multiple copies of *3xP3-RFP* lead to increased RFP fluorescence and ectopic expression of Gal4. Unfortunately, we could not test this because our efforts to remove *3xP3-RFP* using Cre/*loxP* excision were unsuccessful; progeny expressing Cre and *ebony-Gal4* CRISPaint were apparently lethal, whereas we easily generated RFP-free derivatives of *ebony-Gal4* HDR using the same crossing scheme. In contrast, *Mhc-T2A-Gal4* lines #1, #2, and *Myo1a-T2A-Gal4* #1 do not express in the anal pad, and gut and anal pad expression in *wg-T2A-Gal4* #1 likely corresponds to normal *wg* expression in these tissues (Takashima and Murakami 2001) (Figure 4B). In conclusion, these results suggest that Gal4-expressing CRISPaint insertions can capture the endogenous expression pattern of the target gene, albeit with caveats that we demonstrated (also see discussion).

To facilitate the insertion of other sequences into the germ line to generate expression reporters, we generated 10 additional universal donor plasmids (Figure 4F). These include T2A-containing donors with sequence encoding the alternative binary reporters LexGAD, QF2, and split-Gal4, as well as Cas9 nuclease, FLP recombinase, Gal80 repression protein, NanoLuc luminescence reporter, and super-folder GFP. In addition, we generated *pCRISPaint-sfGFP-3xP3-RFP*, which can be used to insert into 3’ coding sequence, generating a C-terminal GFP fusion protein. Finally, we created a single plasmid expressing all three frame-selector sgRNAs (*pCFD5-frame-selectors_0,1,2*). Since indels at the insertion site in the germ line effectively randomizes the coding frame, we reasoned that simultaneous expression of all three sgRNAs would maximize cutting and linearization of a CRISPaint universal donor plasmid.

### Knock-ins targeting uncharacterized small open reading frame (smORF) genes

To further demonstrate that this method can quickly generate knock-in insertions for any gene, we targeted a class of uncharacterized small open reading frame (smORF) genes. smORF genes encode peptides <100 amino acids and have been historically understudied (Couso and Patraquim 2017). For example, these genes may have been missed in mutagenesis and proteomics screens because of their small size. In addition, they frequently lack long introns useful for isolating exon trap insertions (Buszczak *et al.* 2007).

Using sequence alignment tools (BLASTP, NCBI; DIOPT, www.flyrnai.org/diopt) and gene databases (FlyBase; Alliance of Genome Resources, www.alliancegenome.org/) we identified uncharacterized smORF genes that have clear human homologues and targeted them for knock-in using sgRNAs that cut 5’ coding sequence and the *pCRISPaint-T2A-Gal4-3xP3-RFP* donor plasmid. Using the same injection and crossing scheme described above, we isolated 12 insertions in 5 genes as balanced stocks in 1 month (Supplemental Table 6). Diagnostic PCR analysis confirmed that all the insertions integrated in the correct target site (Supplemental Figure 7). These knock-in alleles will be described elsewhere. These results demonstrate that this is a fast method for producing knock-in lines for any gene.

## Discussion

Inserting large DNA elements into the genome by HDR requires a great deal of expertise and labor for the design and construction of donor plasmids. Some groups have developed strategies to improve the efficiency and scale at which homology arms are cloned into donor plasmids (Housden *et al.* 2014; Gratz *et al.* 2015), but the root problem still remains. In this study, we showed that homology-independent insertion can be used in *Drosophila* as an alternative to HDR for certain gene targeting strategies in cultured cells and in vivo. With no donor design and construction, knock-in lines are simpler and faster to obtain (see Table 1), making it easier to target multiple genes in parallel.

**Table 1.** Knock-in timeline comparison in S2R+ cells and in vivo for CRISPaint and HDR.

| S2R+ gene tagging (e.g. mNeonGreen) | Donor design and construction | sgRNA cloning | Transfection -> single cell clones | Molecular characterization | Screening expression |
| --- | --- | --- | --- | --- | --- |
| HDR knock-in | Variable, 1-3 weeks | 3 days | 2 months | 1-2 days (PCR, Gel image) | 1 day, confocal microscopy |
| CRISPaint knock-in | N/A | 3 days | 2 months | 3 days (PCR, Gel image, Sequence) | 1 day, confocal microscopy |

| In vivo KO | Donor design and construction | sgRNA cloning | Injection -> fly stock | Molecular characterization |
| --- | --- | --- | --- | --- |
| HDR knock-in | Variable, 1-3 weeks | 3 days | 1 month | 1-2 days (PCR, Gel image) |
| In/del frameshift | N/A | 3 days | 1 month | 3-7 days (PCR, Gel image, Sequence) |
| CRISPaint knock-in | N/A | 3 days | 1 month | 3 days (PCR, Gel image, Sequence) |

| In vivo gene tagging (e.g. T2A-Gal4) | Donor design and construction | sgRNA cloning | Injection -> fly stock | Molecular characterization | Screening expression |
| --- | --- | --- | --- | --- | --- |
| HDR knock-in | Variable, 1-3 weeks | 3 days | 1 month | 1-2 days (PCR, Gel image) | Optional though recommended:<br>Cross with UAS-GFP, 5-14 days |
| CRISPaint knock-in | N/A | 3 days | 1 month | 3 days (PCR, Gel image, Sequence) | Cross with UAS-GFP, 5-14 days |

To perform homology-independent insertion in *Drosophila*, we adapted the mammalian CRISPaint system (Schmid-Burgk *et al.* 2016) because of its user-friendly design. CRISPaint donor plasmids are “universal” because they lack homology arms, and publicly available collections enable researchers to “mix and match” insert sequences and target genes. Indeed, we showed that *pCRISPaint-mNeonGreen-T2A-PuroR*, originally used in human cells, functions in *Drosophila* S2R+ cells (Figure 1). To linearize donor plasmids in *Drosophila* in three coding frames, we generated plasmids expressing the three CRISPaint frame-selector sgRNAs under the *Drosophila U6* promoter. Similarly, we also generated a plasmid to express all three sgRNAs simultaneously for maximum cutting efficiency. Therefore, to perform a knock-in experiment, only an sgRNA that targets a gene of interest is needed, which are simple to produce by molecular cloning or ordering. In fact, all target gene sgRNAs plasmids used in this study were obtained from the TRiP facility and an ever-growing number of additional sgRNAs are being generated.

Unlike HDR, which can seamlessly insert any DNA into a target genomic location, homology-independent insertion is restricted to specific targeting scenarios. For example, since the entire *pCRISPaint* donor plasmid integrates into the target site, it cannot be used to replace genes, make amino acid changes, or tag proteins at their N-terminus. In addition, potential concatemerization of the insertion (Auer *et al.* 2014; Kumagai *et al.* 2017) prevents its use for strategies where a single copy of the insertion is required, such as artificial exon traps that exploit target gene splicing (Buszczak *et al.* 2007). Despite these limitations, homology-independent insertion is still well suited to generate loss of function alleles by premature truncation, as well as C-terminal gene tagging when a 3’ UTR is included.

Using CRISPaint in the fly germ line, we rapidly generated loss of function alleles by inserting the selectable marker *3xP3-RFP* into target gene coding sequence. This method has several advantages over isolating frameshift alleles using CRISPR-induced indels (Bier *et al.* 2018). First, identifying founder flies with RFP fluorescence is easier and faster than PCR genotyping/sequencing for frameshift indels. Second, insertion into 5’ coding sequence is more likely to disrupt the target gene, whereas frameshift alleles can sometimes retain protein activity (Tuladhar *et al.* 2019). Though, insertions could also have unintended consequences, such as disrupting neighboring gene expression. Third, genetic crosses are simplified by tracking insertion alleles with a fluorescent marker, which could be especially useful in non-*melanogaster* species that do not have balancer chromosomes. Indeed, the combined use of *3xP3-RFP* and *3xP3-GFP* universal donor plasmids (Figure 3H) could facilitate generating double mutant lines. While we did not find evidence for off-target insertions, knock-in lines should be vetted using traditional techniques, including molecular verification of the insertion site, comparing independent insertions, outcrossing to wild type, and performing complementation tests.

Using homology-independent insertion for gene tagging requires screening for properly expressed inserts, due to the unpredictable and error-prone nature of NHEJ. For example, donor plasmids can integrate in two orientations and indels at the insertion site can shift the coding frame. In cultured S2R+ cells, we used antibiotic resistance to efficiently select for mNeonGreen protein fusions (Figure 1D). However, such an assay is not feasible in flies, and so we screened germ line *T2A-Gal4* insertions for expression by crossing with *UAS-GFP* (Figure 3B, Figure 4). This yielded Gal4-expressing lines for *ebony, wg, Mhc*, and *myo1a*. In contrast, we obtained no insertions for *alpthaTub84B, btl, Desat, ap*, and one non-Gal4 expressing insertion each for *FK506-bp2, esg*, and *hh*. This highlights the importance of obtaining multiple independently derived germ line insertions to screen for insert expression. Additional steps could be taken to obtain more insertions per gene, such as increasing the number of injected embryos or reattempting with a different sgRNA. Since our knock-in efficiency is roughly similar to HDR (5-21% (Figure 3C) vs. 5-22% (Gratz *et al.* 2014), 46-88% (Port *et al.* 2015), and 7-42% (Gratz *et al.* 2015)), the limited number of independent germ line insertions may simply be a constraint of embryo injection-based transgenesis.

Sequence analysis of insertion sites in cultured cells and in vivo revealed expected and unexpected results. Single cell cloned S2R+ lines expressing mNeonGreen fusion proteins had insertion sites that were seamless (7/9) or caused an in-frame 3bp deletion (2/9), which is consistent with human cell data (Schmid-Burgk *et al.* 2016). However, all sequenced germ line insertions (20) contained indels at the insertion site, and indels were larger than in S2R+ cells (Supplemental Figure 3). Germ cells are known to differ in their NHEJ mechanisms compared to somatic cells (Preston *et al.* 2006; Ahmed *et al.* 2015), but it is not clear why this would prevent seamless insertions. In addition, 4/6 of Gal4-expressing insertions were out of frame relative to the target gene (Supplemental Figure 3). We speculate that this could result from ribosome frameshifting (Ketteler 2012), an internal ribosome entry site (IRES) (Komar and Hatzoglou 2005), alternative open reading frames (altORFs) (Mouilleron *et al.* 2016), or Gal4 translation initiation from its own start codon. Ultimately, we consider this a fortuitous effect as long as Gal4 is expressed in the correct pattern. Conversely, *wg* #6 is in-frame but does not express Gal4, perhaps due to a mutation of Gal4 sequences. Indeed, splinkerette sequencing results from 3/20 of our RFP+ insertions suggest that portions of the donor plasmid may be deleted during insertion (Supplemental Table 4). These findings again highlight the importance of screening insertions for insert expression.

For gene-tagged cell lines and fly lines generated using homology-independent insertion, potential artifacts that could impact faithful protein fusion or reporter expression. First, indels at the insertion site could introduce cryptic splice sites, delete an important sequence region, or cause non-optimal codons that impact translation of the fusion protein. Second, bacterial sequences in the inserted plasmid may cause transgene silencing or impact neighboring gene expression (Chen *et al.* 2004; Suzuki *et al.* 2016). Though, we note that thousands of transgenic fly lines contain bacterial sequences from phiC31 integration (Perkins *et al.* 2015) with no reports of ill effects. Third, concatemer insertions could affect gene expression in unpredictable ways. Indeed, insertion *ebony-T2A-Gal4 pFP545 #*2 was a head-to-tail concatemer (Supplemental Figure 4) and had ectopic Gal4 expression from the *3xP3* enhancer (Figure 4D,E). Finally, as noted above, the insertion frame may be important for expression, since *wg-T2A-Gal4* #4 line is expressed weakly and in a non-faithful patchy pattern in the wing disc. While these are important considerations, we note that 9/9 of our mNeonGreen tagged S2R+ lines were localized properly, and 5/6 Gal4 expressing lines appear to be faithful reporters of their target genes, with the above exceptions noted.

Some of our S2R+ mNeonGreen protein fusion lines exhibited unusual or unexpected protein localization. In clones D6 and D9, Lamin-mNeonGreen localized to the nuclear envelope, but in D9 this localization was enriched asymmetrically in the direction of the previous plane of cell division. Since both clones had seamless mNeonGreen insertions, we believe the localization difference is caused by the lack of a non-knock-in locus in D9, whereas D6 contained an in-frame 3bp deletion at the non-knock-in locus, likely retaining wild-type function. We saw a similar pattern for clones A3 and A5, both of which had seamless insertions in *Actin5c*, but clone A5 exhibited distinct rod structures. alphaTub84B-mNeonGreen fluorescence and protein levels were low in all cell lines, despite *alphaTub84B* being highly expressed in S2R+ cells (Hu *et al.* 2017). We speculate that the alphaTub84B-mNeonGreen fusion protein is unstable and previous studies in other organisms have highlighted problems with C-terminal tagging of alpha-Tubulin (Carminati and Stearns 1997). Similarly, C-terminal tags can disrupt Lamin and Actin function (Davies *et al.* 2009; Nagasaki *et al.* 2017). Therefore, it is important to consider the existing knowledge of the protein when generating C-terminal protein fusions and examine multiple single cell cloned lines. Importantly, this level of scrutiny is also required when using HDR to produce knock-ins. Further validation steps could be done, such as comparing the tagged protein localization to antibody staining of the untagged wild type protein.

Additional applications of homology-independent insertion in the *Drosophila* germ line can be imagined. For example, new insert sequences could be added to *pCRISPaint-3xP3-RFP* or *pCRISPaint-3xP3-GFP* starting plasmids. In addition, a modified donor plasmid could be constructed that does not integrate the bacterial plasmid backbone, such as providing donor plasmids as mini-circles (Schmid-Burgk *et al.* 2016; Suzuki *et al.* 2016), cutting donor plasmid twice to liberate the insert fragment (Lackner *et al.* 2015; Suzuki *et al.* 2016; Gao *et al.* 2019), or using PCR amplified inserts (Manna *et al.* 2019). Protein tagging could be performed in vivo, but the large indels observed in germ line may limit its use to proteins that have C-terminal tails that can be deleted without altering protein function. Regardless, we generated a donor plasmid for in vivo sfGFP C-terminal fusions (Figure 4F). Finally, we note that most of our germ line donor plasmids contain enzyme restriction sites that can be used to insert genomic homology arms by molecular cloning, making them “dual use” reagents for HDR and homology-independent insertion. Indeed, we used this approach to generate the *ebony-Gal4* HDR allele (see Materials and Methods).

In summary, we demonstrated that homology-independent insertion can be used as an alternative to HDR in *Drosophila*, enabling researchers to rapidly obtain knock-ins without donor design and construction. The most practical application of this approach is to perform C-terminal protein tagging in cultured cells and gene knockout by insertion in vivo. While we obtained in vivo *T2A-Gal4* gene-tagged insertions, it is low efficiency and thus less appealing compared to HDR. However, the techniques required for screening insertions for *T2A-Gal4* are less specialized than those for constructing donor plasmids, making this trade off potentially attractive for those with less molecular biology expertise or who previously failed using HDR. Finally, our in vivo donor plasmids are immediately useable in other arthropod species because of the *3xP3*-fluorescent marker (Berghammer *et al.* 1999), a testament to the modularity of this knock-in system and the benefits of a community of researchers creating and sharing universal donor plasmids.

## Acknowledgements

We thank Jonathan Schmid-Burgk for advice and the *CRISPaint-mNeonGreen* donor plasmid, Claire Hu and the TRiP for sgRNA design and construction, Ben Ewen-Campen for valuable comments on the manuscript, Stephanie Mohr, Oguz Kanca, and Hugo Bellen for helpful discussions, Raghuvir Viswanatha for the *Cas9-T2A-EGFP* template sequence, Rich Binari for help with mounting embryos on slides, and Cathryn Murphy for general assistance. We also thank the HMS MicRoN (Microscopy Resources on the North Quad) Core and the HMS Department of Immunology’s Flow Cytometry Facility for technical assistance. J.A.B. was supported by the Damon Runyon Foundation. This work was supported by NIH grants R01GM084947, R01GM067761, R24OD019847. N.P. is an investigator of the Howard Hughes Medical Institute.

---

**Supplemental Figure 1.**
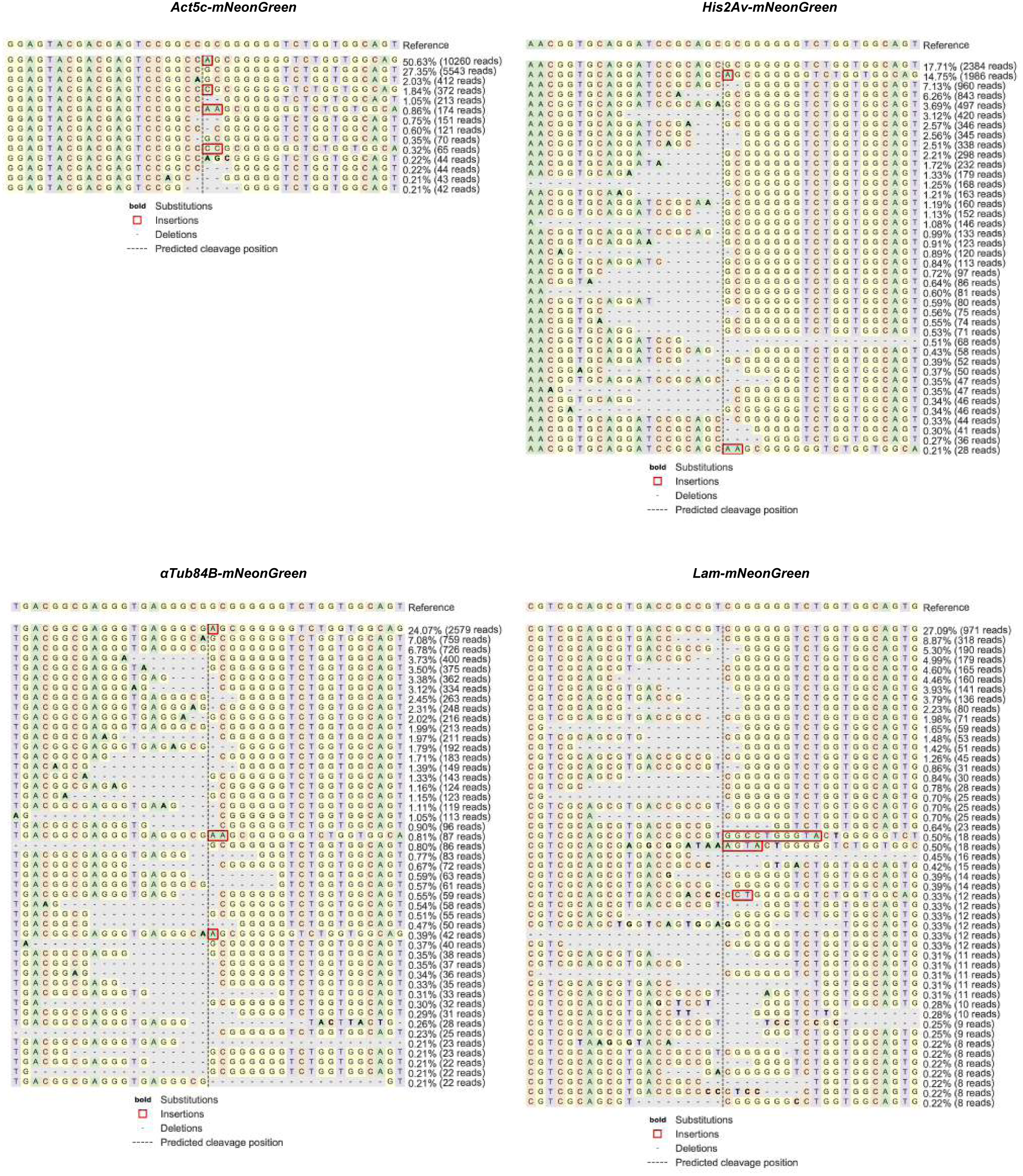
Next-generation sequencing analysis of mNeonGreen insertion sites in transfected S2R+ cells using CRISPresso2

**Supplemental Figure 2.**
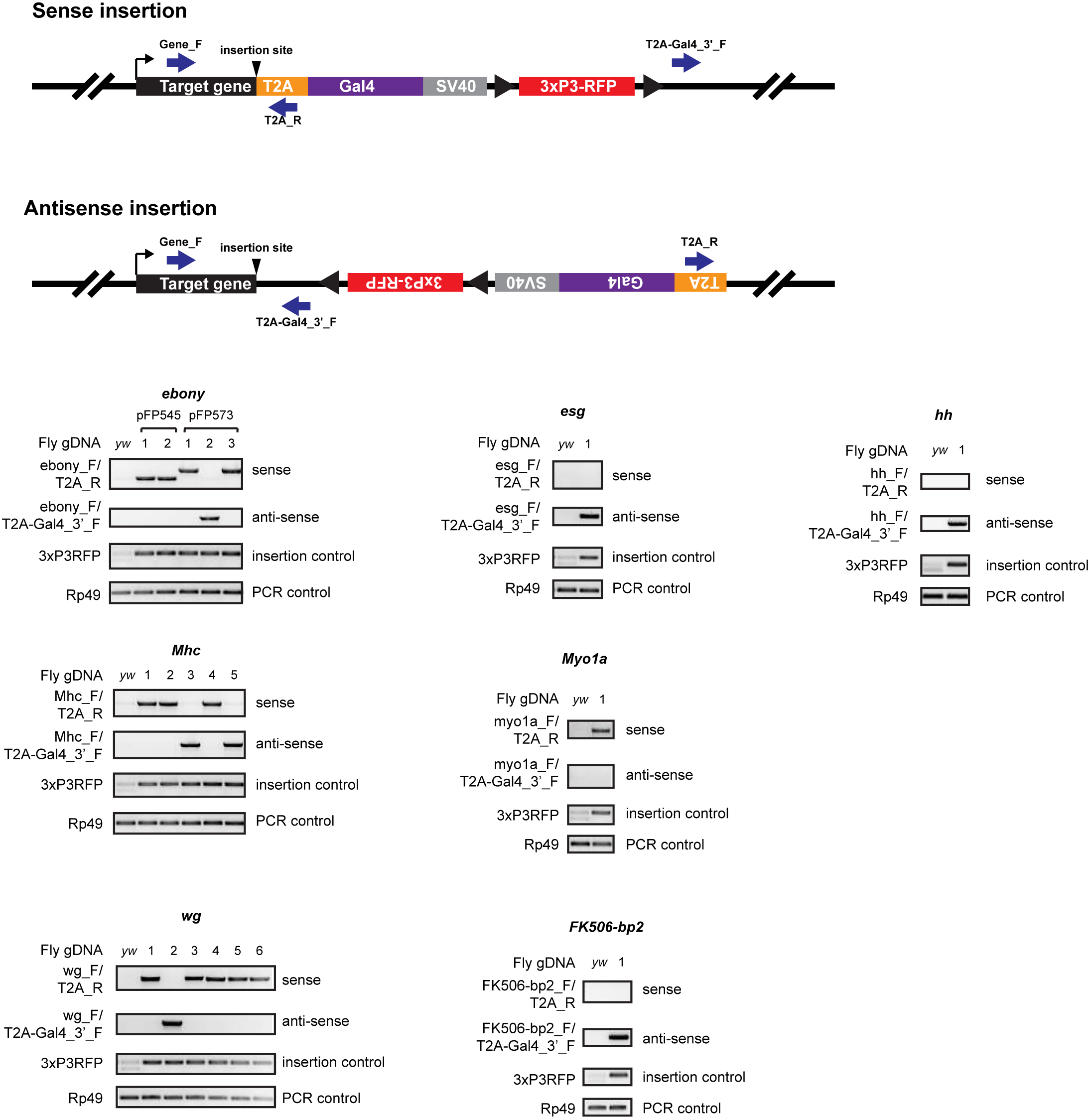
Diagnostic PCR of *CRISPaint-T2A-Gal4* insertions in RFP+ fly lines to confirm their insertion site and orientation.

**Supplemental Figure 3.**
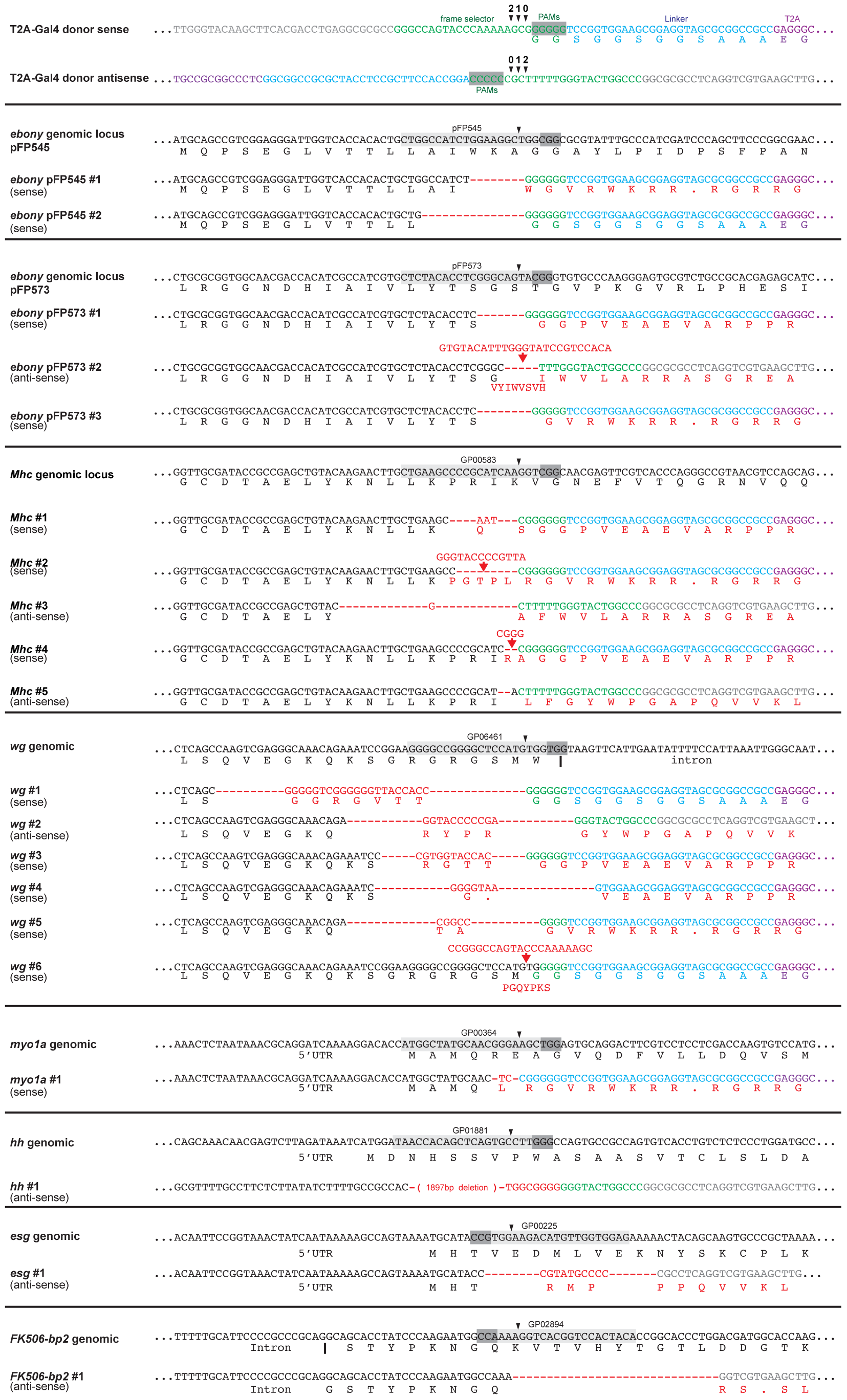
Sequence structure of *CRISPaint-T2A-Gal4* insertions in RFP+ fly lines

**Supplemental Figure 4.**
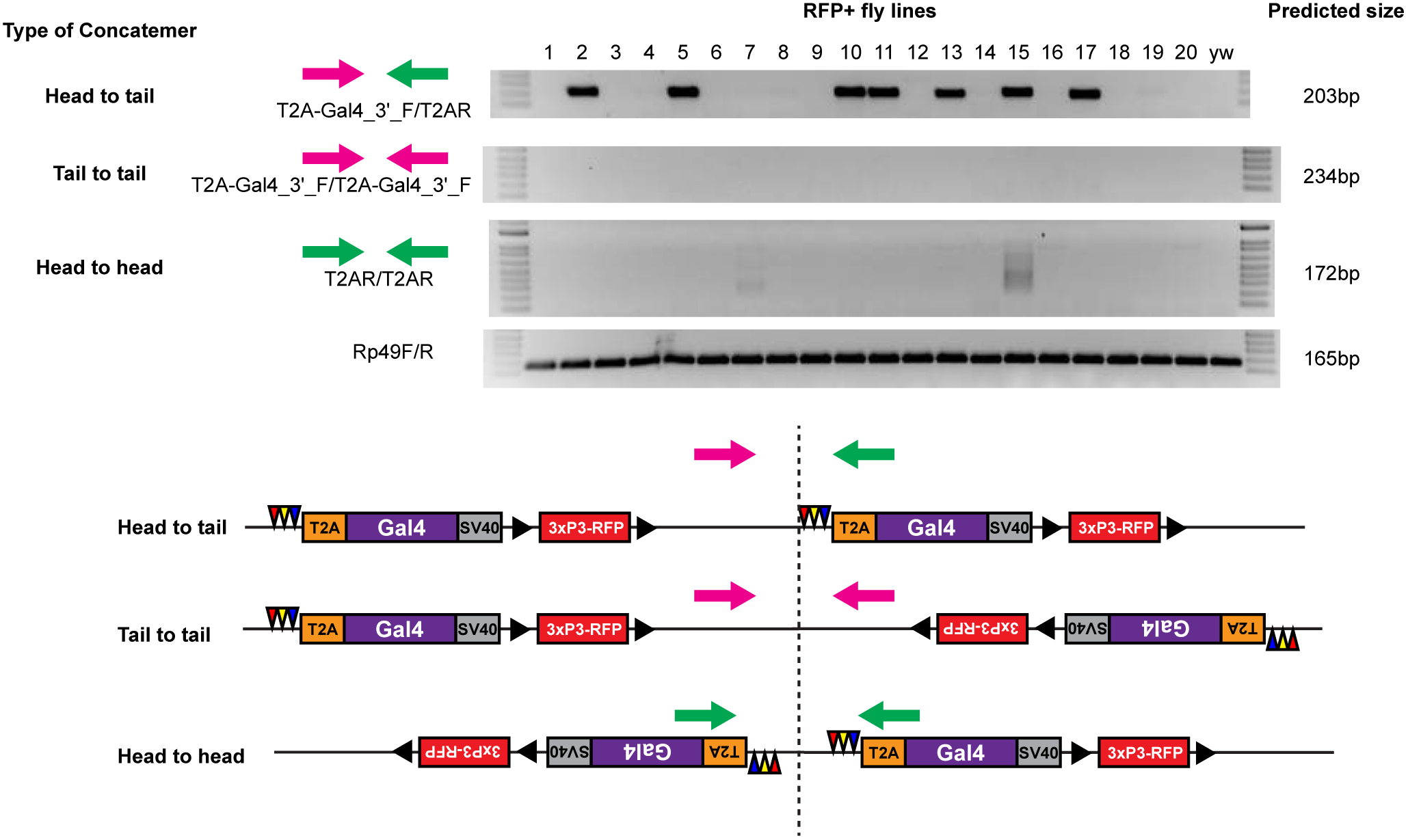
Diagnostic PCR of *CRISPaint-T2A-Gal4* insertions in RFP+ fly lines to detect concatemers

**Supplemental Figure 5.**
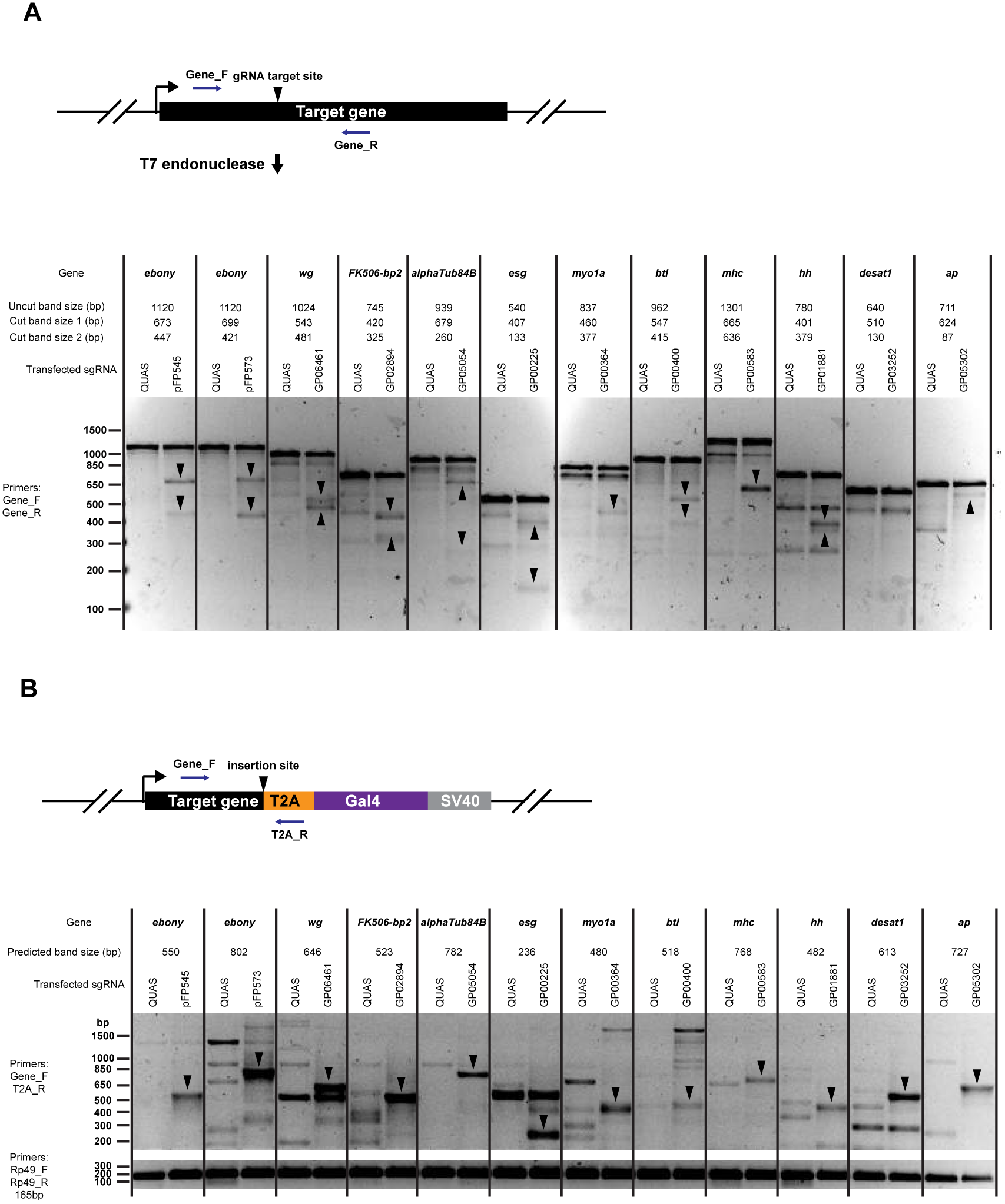
Cutting efficiency of 12 sgRNAs in transfected S2R+ cells. (**A**) T7 endonuclease assay. (**B**) Diagnostic PCR to detect for presence of sense orientation *CRISPaint-T2A-Gal4* insertion events.

**Supplemental Figure 6.**
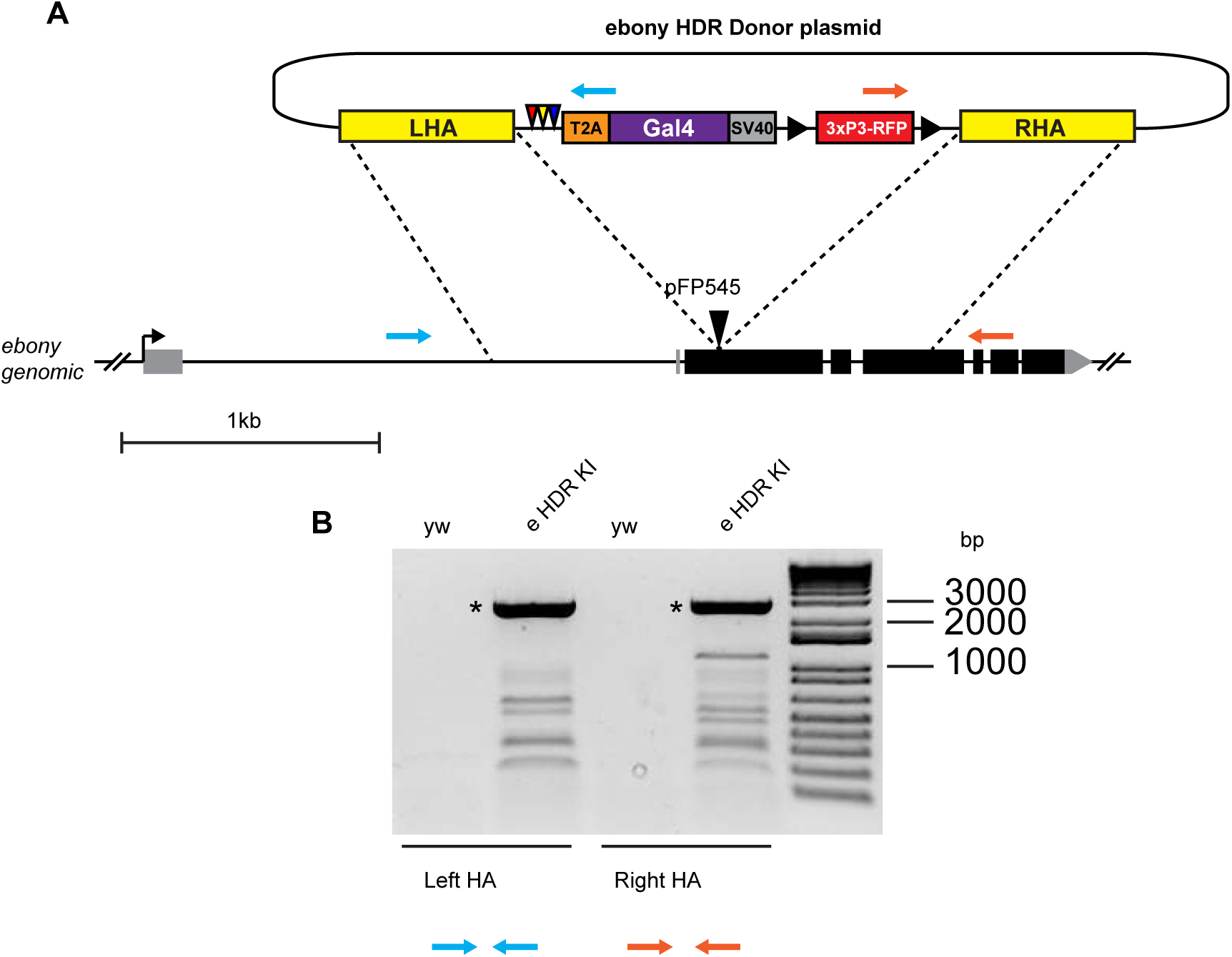
Construction of *ebony-Gal4* HDR insertion. (**A**) Targeting strategy of *ebony* HDR donor plasmid and *ebony* genomic locus. (**B**) Diagnostic PCR of *ebony-Gal4* HDR insertion. Asterisks indicate correct size band.

**Supplemental Figure 7.**
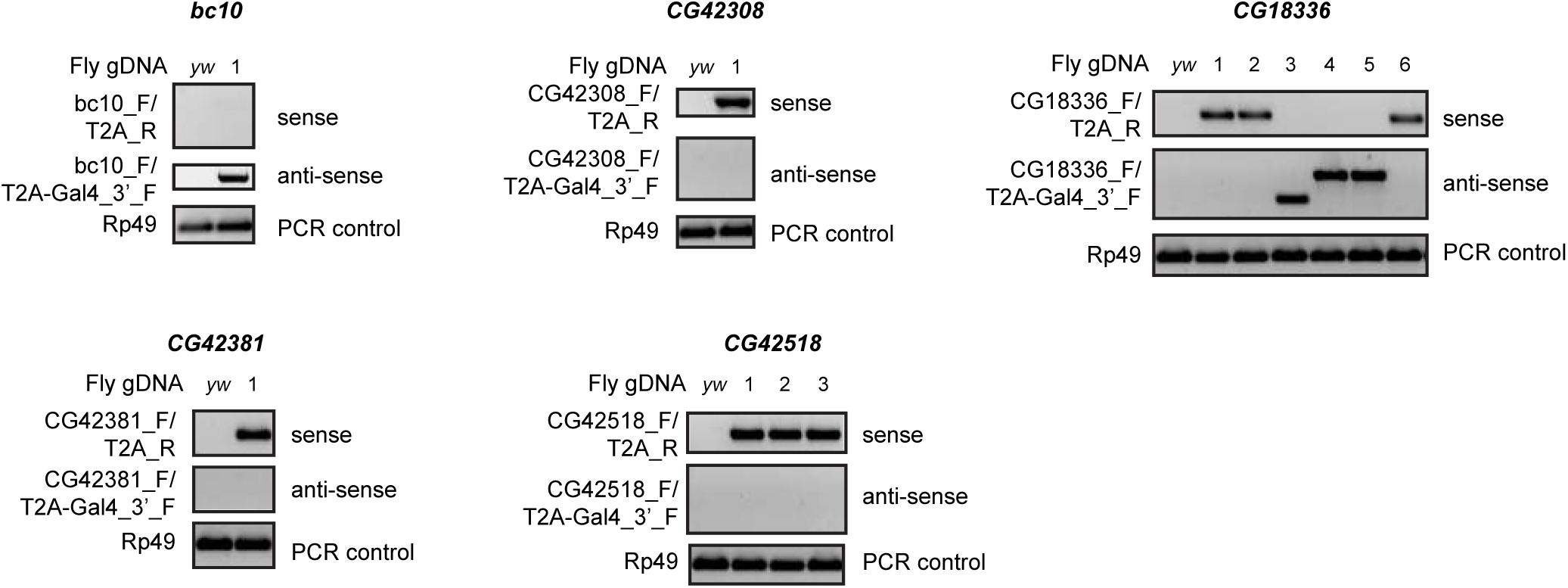
Diagnostic PCR of CRISPaint-T2A-Gal4 insertions in smORF genes.

**Supplemental Table 1.**
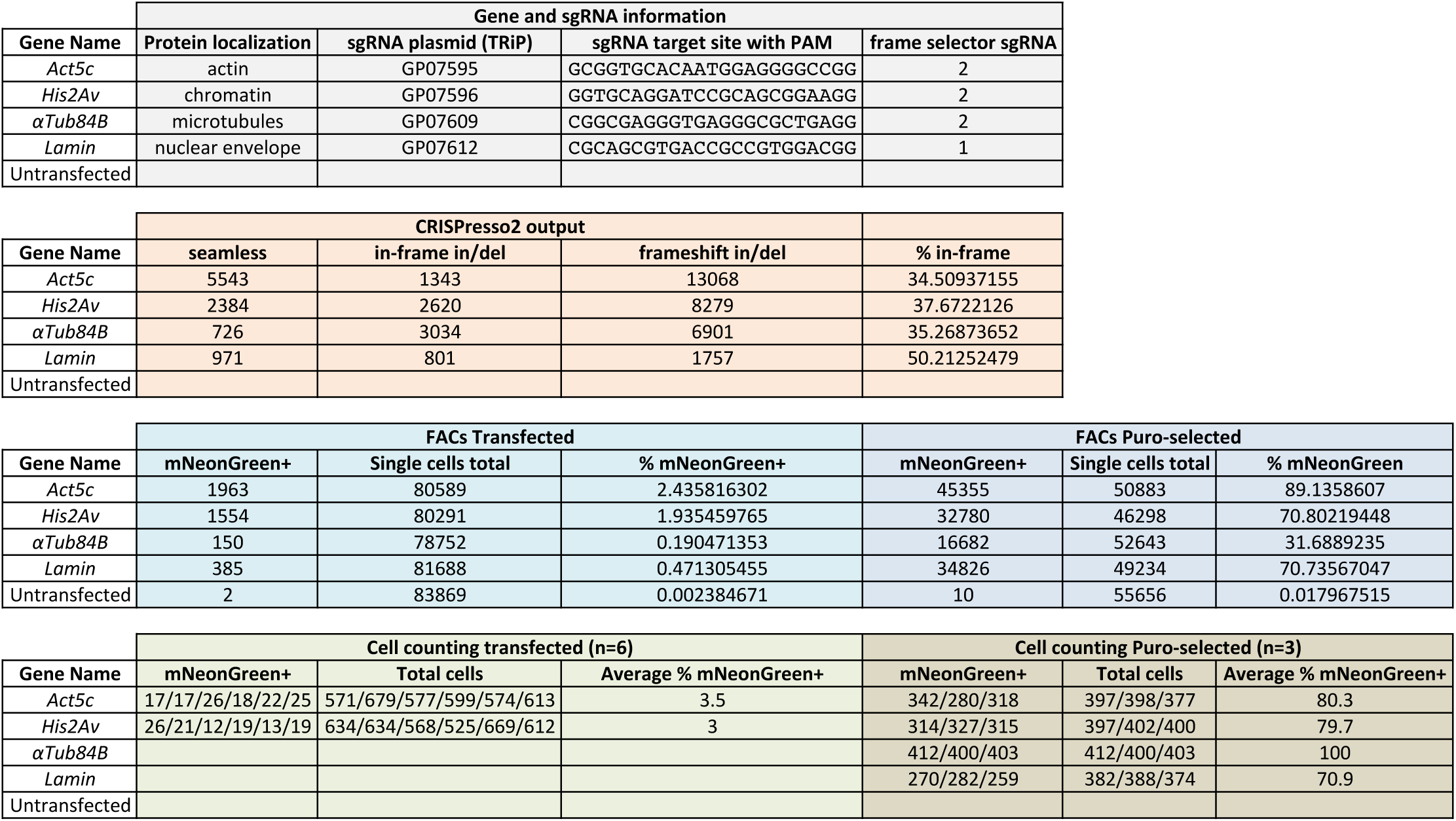
Quantification of CRISPaint-mNeonGreen insertion events in transfected and puro-selected S2R+ cells by CRISPresso2, FACs, and cell counting of confocal images.

**Supplemental Table 2.**
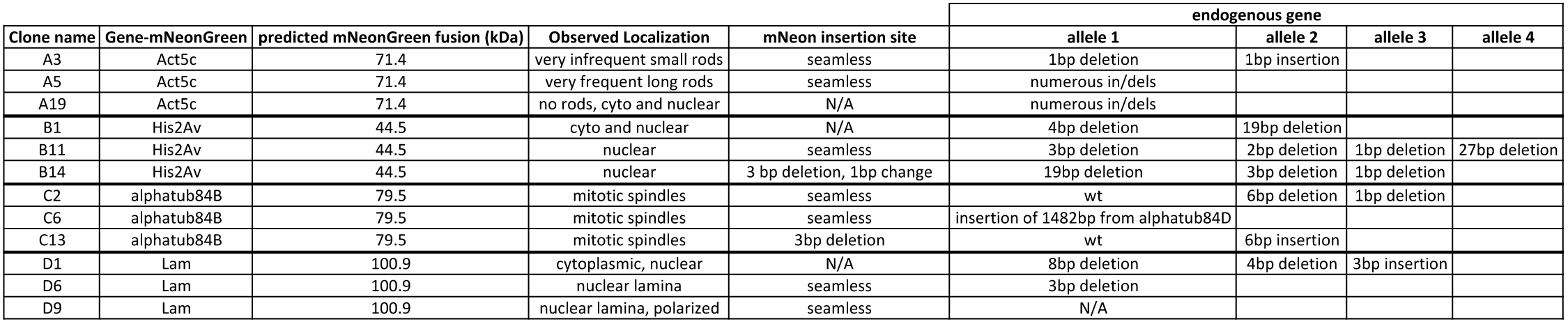
Molecular characterization of single cell cloned mNeonGreen-expressing S2R+ lines.

| Clone name | Gene-mNeonGreen | predicted mNeonGreen fusion (kDa) | Observed Localization | mNeon insertion site | endogenous gene |  |  |  |
| --- | --- | --- | --- | --- | --- | --- | --- | --- |
|  |  |  |  |  | allele 1 | allele 2 | allele 3 | allele 4 |
| A3 | Act5c | 71.4 | very infrequent small rods | seamless | 1bp deletion | 1bp insertion |  |  |
| A5 | Act5c | 71.4 | very frequent long rods | seamless | numerous in/dels |  |  |  |
| A19 | Act5c | 71.4 | no rods, cyto and nuclear | N/A | numerous in/dels |  |  |  |
| B1 | His2Av | 44.5 | cyto and nuclear | N/A | 4bp deletion | 19bp deletion |  |  |
| B11 | His2Av | 44.5 | nuclear | seamless | 3bp deletion | 2bp deletion | 1bp deletion | 27bp deletion |
| B14 | His2Av | 44.5 | nuclear | 3 bp deletion, 1bp change | 19bp deletion | 3bp deletion | 1bp deletion |  |
| C2 | alphatub84B | 79.5 | mitotic spindles | seamless | wt | 6bp deletion | 1bp deletion |  |
| C6 | alphatub84B | 79.5 | mitotic spindles | seamless | insertion of 1482bp from alphatub84D |  |  |  |
| C13 | alphatub84B | 79.5 | mitotic spindles | 3bp deletion | wt | 6bp insertion |  |  |
| D1 | Lam | 100.9 | cytoplasmic, nuclear | N/A | 8bp deletion | 4bp deletion | 3bp insertion |  |
| D6 | Lam | 100.9 | nuclear lamina | seamless | 3bp deletion |  |  |  |
| D9 | Lam | 100.9 | nuclear lamina, polarized | seamless | N/A |  |  |  |

**Supplemental Table 3.**
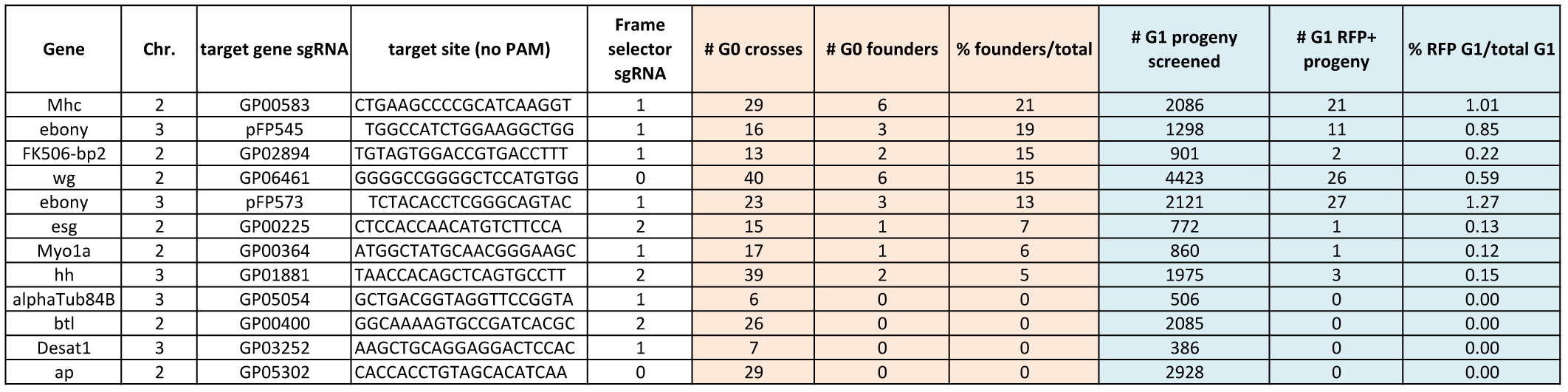
Germ line knock-in efficiency of *CRISPaint-T2A-Gal4*.

**Supplemental Table 4.** Molecular and phenotypic characterization of 20 RFP+ fly strains, each carrying a distinct *CRISPaint-T2A-Gal4* insertion.

| # | Gene targeted | sgRNA | G0 cross # | Head-tail insertion by PCR? | Splinkerette 5' (PCR Purification) | Splinkerette 5' (Gel Purification) | Splinkerette 3' (PCR Purification) | Splinkerette 3' (Gel Purification) | Insert orientation | Insert frame | Insertion site sequence | Gal4 expression | Homozygous phenotype | Complementation test |
| --- | --- | --- | --- | --- | --- | --- | --- | --- | --- | --- | --- | --- | --- | --- |
| 1 | ebony | pFP545 | 1 | no | ebony 5' insertion site | no bands | ebony 3' insertion site | ebony 3' insertion site | Sense | Out | 8bp del | none | dark cuticle pigment | dark cuticle pigment over e1 on TM3 |
| 2 | ebony | pFP545 | 2 | yes | ebony 3' insertion site | Band 1: ebony 3' insertion site<br>Band 2: 3' end of donor plasmid | 5' end of donor plasmid | 5' end of donor plasmid | Sense | In | 15bp del | anterior and posterior tracheae in larvae, whole body cuticle in pupae and adult | lethal | dark cuticle pigment over e1 on TM3 |
| 3 | ebony | pFP573 | 1 | no | internal donor plasmid | internal donor plasmid | vermillion | vermillion | Sense | Out | 7bp del | none | dark cuticle pigment | dark cuticle pigment over e1 on TM3 |
| 4 | ebony | pFP573 | 2 | no | ebony 3' insertion site | ebony 3' insertion site | vermillion | vermillion | Antisense | N/A | 5bp del, 25bp ins | none | dark cuticle pigment | dark cuticle pigment over e1 on TM3 |
| 5 | ebony | pFP573 | 3 | yes | 3' end of donor plasmid | 3' end of donor plasmid | vermillion | bad sequence data | Sense | Out | 8bp deletion | none | dark cuticle pigment | dark cuticle pigment over e1 on TM3 |
| 6 | hh | GP01881 | 1 | no | bad sequence data | no bands | vermillion | Band 1: hh 5' insertion site<br>Band 2: hh 3' insertion site<br>Band 3: vermillion | Antisense | N/A | 1897bp del, 8bp ins | none | lethal | fail to complement hh[AC] and Df(3R)ED5296 |
| 7 | Mhc | GP00583 | 1 | no | Mhc 5' insertion site | no bands | vermillion | Band 1: Mhc 5' insertion site<br>Band 2: vermillion | Sense | Out | 10bp del, 3bp ins | muscle | lethal | fail to complement P(lacW)Mhck10423 and Df(2L)H20 |
| 8 | Mhc | GP00583 | 2 | no | Mhc 5' insertion site | no bands | bad sequence data | bad sequence data | Sense | Out | 9bp del, 13bp ins | muscle | lethal | fail to complement P(lacW)Mhck10423 and Df(2L)H20 |
| 9 | Mhc | GP00583 | 3 | no | Mhc 3' insertion site | Mhc 3' insertion site | bad sequence data | bad sequence data | Antisense | N/A | 27bp del, 1bp ins | none | lethal | fail to complement P(lacW)Mhck10423 and Df(2L)H20 |
| 10 | Mhc | GP00583 | 4 | yes | Mhc 3' insertion site | Band 1: 3' end of donor plasmid<br>Band 2: 3' end of donor plasmid | bad sequence data | Band 1: 5' end of donor plasmid<br>Band 2: internal donor plasmid | Sense | Out | 2bp del, 4bp ins | none | lethal | fail to complement P(lacW)Mhck10423 and Df(2L)H20 |
| 11 | Mhc | GP00583 | 5 | yes | Mhc 3' insertion site | Band 1: Mhc 3' insertion site<br>Band 2: 3' end of donor plasmid | 5' end of donor plasmid | 5' end of donor plasmid | Antisense | N/A | 2bp deletion | none | lethal | fail to complement P(lacW)Mhck10423 and Df(2L)H20 |
| 12 | Myo1a | GP00364 | 1 | no | Myo1a 5' insertion site | Myo1a, 5' insertion site | internal donor plasmid | internal donor plasmid | Sense | Out | 4bp del, 2bp ins | gut | viable | N/A |
| 13 | esg | GP00225 | 1 | yes | esg, 3' insertion site | Band 1: esg, 3' insertion site<br>Band 2: bad sequence data | vermillion | Band 1: 5' end of donor plasmid<br>Band 2: vermillion | Antisense | N/A | 25bp del, 10bp | none | lethal | fail to complement P(enG)esg[G66] |
| 14 | FK506-bp2 | GP02894 | 1 | no | FK506-bp2 3' insertion site | FK506-bp2 3' insertion site | FK506-bp2 5' insertion site | Band 1: FK506-bp2 5' insertion site<br>Band 2: vermillion | Antisense | N/A | 30bp del | none | viable | N/A |
| 15 | wg | GP06461 | 1 | yes | wg 3' insertion site | Band 1: wg 3' insertion site<br>Band 2: 3' end of donor plasmid | vermillion | Band 1: wg 5' insertion site<br>Band 2: 5' end of donor plasmid | Sense | In | 45bp del, 21bp ins | wingless, strong expression | lethal | fail to complement wg[I-17], wg[I-8], and Df(2L)BSC291 |
| 16 | wg | GP06461 | 2 | no | 3' end of donor plasmid | 3' plasmid | wg 5' insertion site | 5' end of donor plasmid | Antisense | N/A | 33bp del, 11bp ins | none | lethal | fail to complement wg[I-17], wg[I-8], and Df(2L)BSC291 |
| 17 | wg | GP06461 | 3 | yes | wg 5' insertion site | Band 1: wg 5' insertion site<br>Band 2: wg 3' insertion site<br>Band 3: 3' end of donor plasmid | vermillion | wg 3' insertion site | Sense | Out | 21bp del, 11bp ins | none | lethal | fail to complement wg[I-17], wg[I-8], and Df(2L)BSC291 |
| 18 | wg | GP06461 | 4 | no | wg 5' insertion site | Band 1: wg 5' insertion site<br>Band 2: 3' end of donor plasmid | wg 3' insertion site | Band 1: wg 3' insertion site<br>Band 2: attB sequence? | Sense | Out | 32bp del, 7bp ins | wingless, weak expression | lethal | fail to complement wg[I-17], wg[I-8], and Df(2L)BSC291 |
| 19 | wg | GP06461 | 5 | no | wg 5' insertion site | wg 5' insertion site | wg 3' insertion site | vermillion | Sense | Out | 28bp del, 5bp ins | none | lethal | fail to complement wg[I-17], wg[I-8], and Df(2L)BSC291 |
| 20 | wg | GP06461 | 6 | no | wg 5' insertion site | Band 1: wg 5' insertion site<br>Band 2: wg 3' insertion site | vermillion | no band | Sense | In | 21bp insertion | none | lethal | fail to complement wg[I-17], wg[I-8], and Df(2L)BSC291 |

**Supplemental Table 5.**
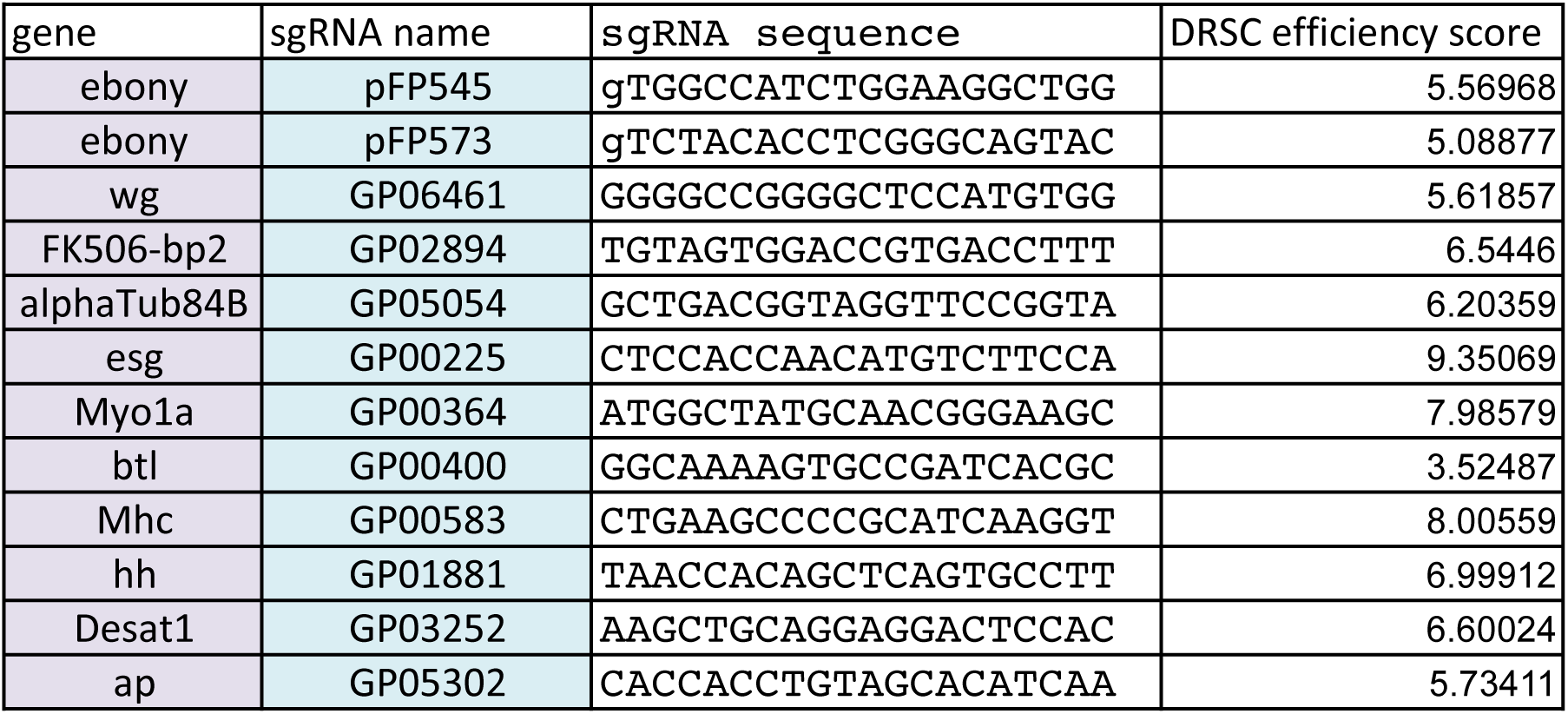
Efficiency scores for 12 sgRNAs used in germ line knock-ins.

**Supplemental Table 6.**
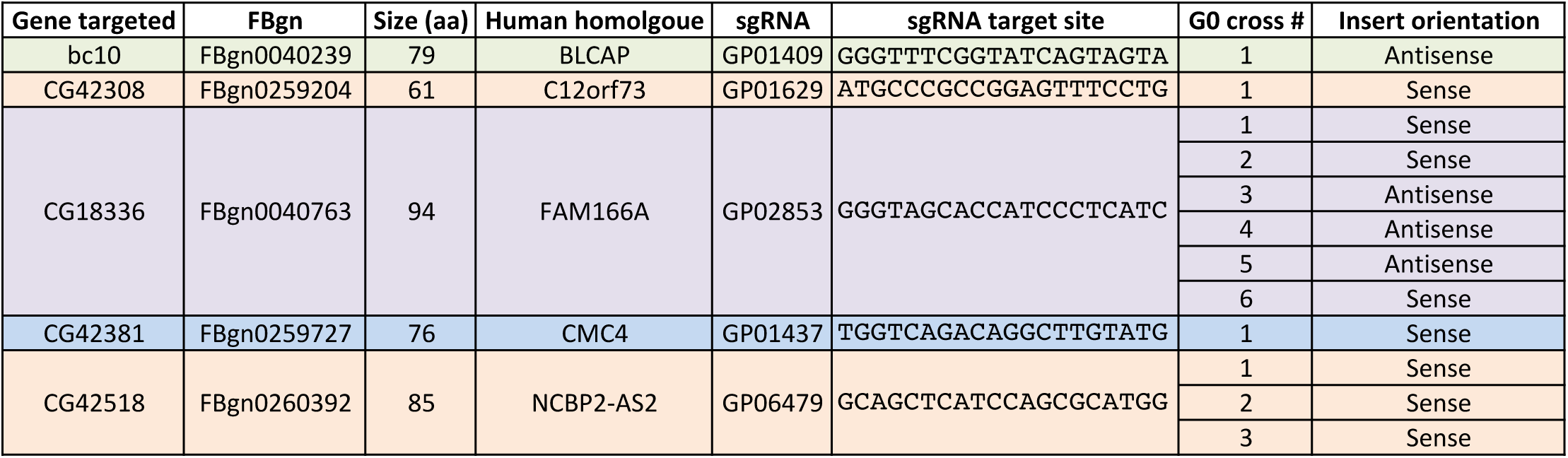
smORF genes targeted for knock-in with pCRISPaint-T2A-Gal4-3xP3-RFP

**Supplemental Table 7.**
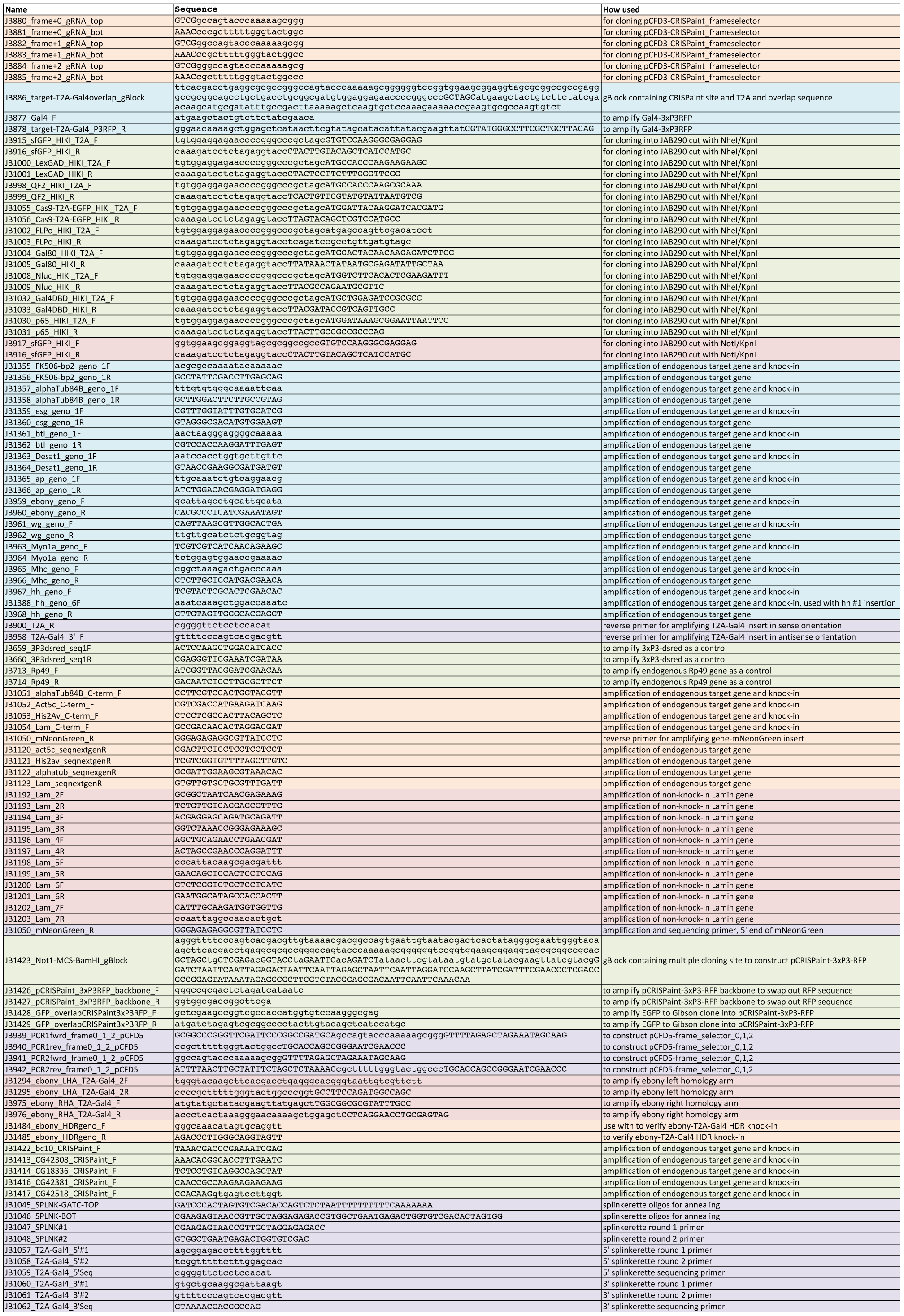
Oligo and dsDNA sequences.

